# Deep learning cell type classification using nuclear DNA patterns

**DOI:** 10.64898/2026.04.28.721280

**Authors:** Kaito Sugimoto, Hiromitsu Tanaka, Tetsuichiro Saito

## Abstract

Multicellular organisms comprise various types of cells, which are characterized by gene expression through interactions between chromosomal DNA and nuclear proteins. Many cutting-edge methods have been developed to reveal the three-dimensional organization of chromosomes. The detailed analyses of whole chromosomes have begun to uncover structural features specific to several cell types. Here, we show that cell types are instantly and highly accurately classified using conventional DNA staining and a convolutional neural network (CNN). A high-resolution single slice image of the nucleus is sufficient for the accurate classification of both live and fixed cells, including neurons and non-neural cells. These findings suggest that there may be cell-type-specific features decipherable by deep learning in a thin two-dimensional slice of the nucleus.

## INTRODUCTION

Many cell types have been found since the microscope was developed. Initially, cell types were classified based on their morphologies. ^1^ Then, molecular markers such as proteins and RNAs that are specifically expressed in a cell type of interest have allowed for its precise classification. Their expression is regulated by many nuclear proteins, including transcription factors and various types of histones, the binding sites of which are mapped by chromatin immunoprecipitation. Chromosomal loci in spatial proximity to each other are analyzed by chromatin conformation capture techniques, such as high-throughput chromosome conformation capture. ^2,3^ The positions of chromosomal loci, RNAs and proteins are directly visualized by in situ hybridization such as DNA seqFISH+ and immunostaining. ^4^ Each chromosome occupies a territory, albeit partially overlapping with other chromosomes, and is separated into structures, such as the A and B compartments, which are enriched for transcriptionally active and inactive genes, respectively, and further subdivided into topologically associating domains and chromatin loops. Whereas bulk analyses using a population of cells suggest the existence of cell-type-specific structural features in chromosomes of some cell types, single-cell analyses have revealed cell-to-cell heterogeneity even within the same cell type. ^5^

On the other hand, CNNs, a type of deep-learning algorithm, have been widely used for image recognition and classification, ^6–9^ and serve as diagnostic tools in the clinic. ^10^ Moreover, CNNs have been applied to predict primary origins for cancers using tissue sections ^11^ and to analyze cellular states, such as cell cycle phases, virus infection and senescence, using DNA-stained images of nuclei. ^12–14^ Here, we present a simple yet powerful approach combining conventional DNA staining with CNNs for the classification of cell types. After staining nuclei with a fluorescent dye, Hoechst 33342 (Hoechst) or 4’,6-diamidino-2-phenylindole (DAPI), we obtain their high-resolution images using a confocal microscope that is combined with a 160× objective lens for a total internal reflection fluorescence microscope. Hoechst and DAPI preferentially bind adenine-thymine (AT)-rich regions, ^15,16^ with higher affinity to heterochromatic regions, ^17,18^ thereby showing variable intensities of DNA staining. CNN models trained on live-cell nuclei images show accurate prediction for live neurons and non-neural cells. Similarly, several cell types in fixed tissues are classified using different CNN models trained on cell nuclei of the tissues. Moreover, training in which four representative subtypes of neurons are categorized into a single unified group of neurons leads to accurate sorting of other untrained subtypes of neurons into the group. Despite huge varieties of neuronal subtypes, ^19^ this finding suggests that there may be common nuclear features among neurons.

## RESULTS

### Classification of live cultured cells

We sought to determine whether live cultured cells might be classified based on the features of their own nuclei. Four well-characterized mouse cells were cultured: P19 embryonal carcinoma (P19) cells, ^20^ neurons, astrocytes and NIH3T3 fibroblasts (Figure 1A). ^21^ The neurons were generated from P19 cells *in vitro.* ^22^ Cell nuclei were stained with Hoechst, which is commonly used for live cells. ^23^ To capture high-resolution images of the nucleus, we used the 160× objective lens (Figure S1) and cropped interphase nuclei (Figure 1B). The nuclear area of neurons was significantly smaller than those of other cell types (Figure 1C). We applied a random forest (RF) algorithm, ^24,25^ which has been effectively used as a classifier. We trained an RF based on twelve features of the nucleus, including nuclear area and circularity etc., with stratified five-fold cross-validation to ensure stable performance, using nuclear images from the four cell types (P19 cells, *n =* 1001; neurons, *n =* 1001; astrocytes, *n =* 947; NIH3T3 fibroblasts, *n =* 1007) (Figure S2). The nuclear area seems to be the most important for prediction. The evaluation using independent external test set (*n* = 200 for each cell type) suggests that the model might be able to predict neurons with high accuracy but is not effective at classifying other cell types (Figure 1D).

**Figure 1.**
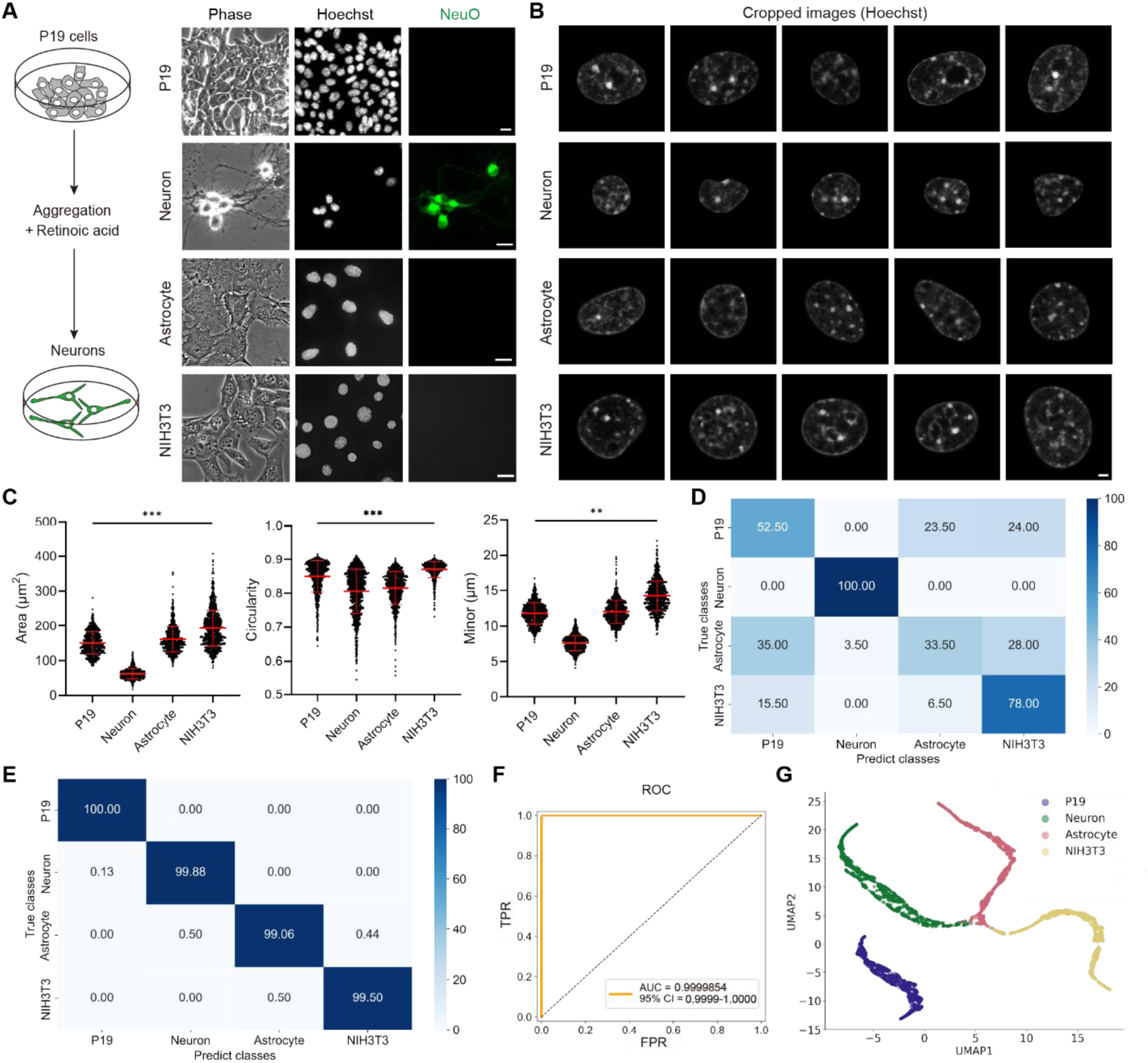
Cell type classification of live cultured cells. (A) Schematic of neuronal differentiation from P19 cells and representative images of P19 cells, neurons, astrocytes and NIH3T3 fibroblasts (Phase contrast, Hoechst and NeuO), after staining with Hoechst and a neuron-selective probe, NeuO. Scale bars, 20 µm. (B) Representative images of cropped nuclei. Scale bar, 2 µm. (C) Quantification of the top three nuclear features (Area, Circularity and Minor), ranked by importance in an RF model. Mean and s.d. are shown. Statistical significance was assessed by Welch’s ANOVA with post-hoc Games-Howell’s test. In Area and Circularity, \*\*\**p* < 0.001 for all pairwise comparisons. In Minor, \*\**p* < 0.01 for the comparison between P19 cells and astrocytes, and *p* < 0.001 for the other comparisons. (D) Confusion matrix of RF model predictions on an external test set. Each value represents the average percentage of correct classification or misclassification for each true class across the five models. (E) Confusion matrix of CNN model predictions on an external test set. (F) ROC curve and AUC for a representative model (model 6 in Table S1). TPR, true positive rate; FPR, false positive rate. (G) UMAP representation of nuclear features learned by the model 6.

Then we employed a CNN, which includes several convolution, max pooling and dropout layers (Figure S3A). Fully connected (FC) layers are applied after flattening feature maps. During training, the cropped images were augmented by random rotation, resizing and brightness adjustment. We trained eight separate CNN models on these images using stratified eight-fold cross-validation (P19 cells, *n =* 15015; neurons, *n =* 15015; astrocytes, *n =* 14205; NIH3T3 fibroblasts, *n =* 15105). Across the eight models, key performance metrics for classifying each cell type, including precision, recall, F1-score and the area under the curve (AUC) of the receiver operating characteristic (ROC), were highly consistent, indicating stable model performance (Table S1). A representative model achieved near-perfect accuracy on the validation set, classifying the four cell types with the accuracy of 99.60 ± 0.50% (mean ± s.d.) (Figure S3B). The validation accuracy increased along with the training accuracy over epochs, and the minimal gap between the two curves suggests that overfitting was prevented during the training (Figure S3C). Similarly, the loss curves for both training and validation sets converged steadily, showing the efficiency of the learning process.

On an external test set (*n* = 200 for each cell type), the average accuracy across the eight models reached 99.61 ± 0.12% (Figure 1E). The AUC was close to 1.000 (Figure 1F).

Uniform manifold approximation and projection (UMAP) ^26^ analysis based on the parameters in the last FC layer of the model revealed clearly separated clusters corresponding to the four cell types (Figure 1G). These results suggest that the CNN models, hereafter designated as live cultured cell (LCC) models, accurately distinguish those four cell types.

Next, we examined whether neurons derived from mouse embryonic stem (ES) cells ^27^ would be also predicted by the LCC models (Figures S4A and S4B). On average, the eight models sorted them into the neuron group with 99.25 ± 0.38% accuracy (Figure S4C). UMAP analysis showed that neurons derived from P19 and ES cells belonged to the same cluster (Figure S4D), suggesting that there might be some neuron-specific nuclear features detectable by deep learning.

### Classification of fixed tissue cells

Next, we tested whether the LCC models could classify cell types in tissues. To determine those cell types, we performed immunostaining after fixing embryonic day 13.5 (E 13.5) mouse cerebral cortex (CTX). Cell nuclei were stained with DAPI, because DAPI is frequently used for fixed cells. ^23^ In the embryonic CTX, neural progenitor cells (NPCs) were identified with their marker, SOX2, a transcription factor essential for NPC maintenance, ^28^ whereas pyramidal neurons were identified with a neuronal marker, TBR1, a transcription factor expressed in several types of neurons ^29^ (Figures 2A, 2B and 2D). At this stage, NPCs and neurons are clearly separated in two distinct layers. Cells undergoing DNA replication were excluded from the datasets, because their nuclear structure was drastically changing. Those cells were identified using labeling with 5-ethynyl-2’-deoxyuridine (EdU) (Figure 2B). The LCC models misclassified NPCs as neurons, although they correctly classified neurons (Figure S5A). We also tested NPCs and neurons that were fixed and stained with Hoechst (Figure S5B). Similarly, the models misclassified the NPCs as neurons (Figure S5C). These results suggest that the LCC models are not capable of accurately classifying cell types in fixed tissues, whether they were stained with DAPI or Hoechst.

**Figure 2.**
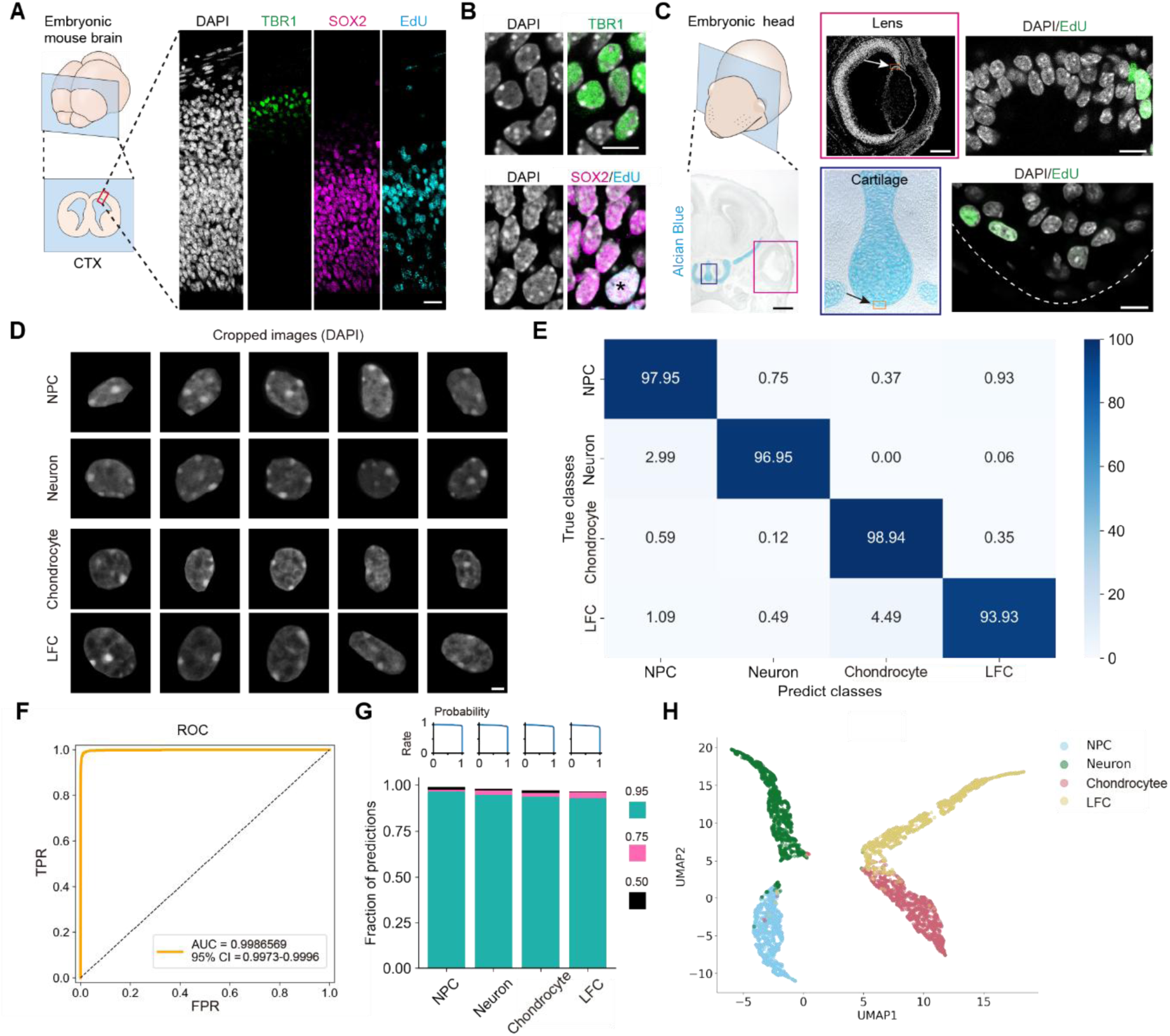
Cell type classification of fixed tissue cells. (A) E13.5 CTX (left, schematic drawing; right, representative immunostaining images). Scale bar, 50 µm. (B) High-magnified images of CTX. Asterisk indicates a SOX2^+^EdU^+^ cell. Scale bar, 10 µm. (C) E15.5 mouse head coronal sections. Left: Schematic drawing and Alcian Blue staining image showing the nasal cartilage (blue box) and lens (magenta box). Scale bar, 500 μm. Middle: Magnified images of the boxes. Scale bar, 200 μm. Right: Magnified images of boxes indicated by arrows in the middle panel. Scale bars, 10 μm. (D) Representative images of cropped nuclei. Scale bar, 2 µm. (E) Confusion matrix of predictions on an external test set. (F) ROC curve and AUC for a representative model (model 7 in Table S2). (G) Top, the fraction (*y* axis) that was correctly classified at or above a certain confidence (probability) threshold (*x* axis) for each cell type. Bottom, the fraction of each cell type that was correctly classified with a confidence score above certain thresholds (>0.95, green; >0.75 and ≤0.95, magenta; >0.50 and ≤0.75, black). (H) UMAP representation of nuclear features learned by the model 7.

Thus, we built new CNN models on cells in fixed tissues, adding two more well-characterized cell types: chondrocytes and lens fiber cells (LFCs) in the embryonic mouse head (Figures 2C and 2D) (NPCs, *n =* 18120; neurons, *n =* 18150; chondrocytes, *n =* 18480; LFCs, *n =* 18120). Chondrocytes in the nasal cartilage ^30^ and LFCs in the peri-central region of the lens, ^31,32^ were identified by their positions, as the morphologies of the cartilage and lens were clear. We performed eight-fold cross-validation using the same CNN architecture as that used for live cells. The eight models showed high and consistent performance, achieving an average accuracy of 96.23 ± 0.48% on validation sets (Figure S5D and Table S2). We then tested the models on an external test set (NPCs, *n =* 201; neurons, *n =* 201; chondrocytes, *n =* 213; LFCs, *n =* 206). The mean accuracy across the eight models reached 96.95 ± 0.38% (Figure 2E). The AUC was close to 1.000 (Figure 2F). Most predictions were made with a high confidence (probability) score above 0.95, indicating general reliability (Figure 2G).

UMAP analysis revealed clear separation of the four cell types (Figure 2H). These results demonstrate the accurate cell type classification in fixed tissues using these CNN models, hereafter designated as fixed tissue cell (FTC) models.

We examined the FTC models on neurons and NPCs that were fixed and stained with Hoechst instead of DAPI (Figure S5B). The FTC models classified NPCs and neurons with average accuracies of 99.38 ± 0.69% and 95.27 ± 2.47%, respectively (Figure S5F), which were comparable to the performance on DAPI-stained cells (Figure 2E), possibly reflecting that DAPI and Hoechst may make similar staining patterns by preferentially binding to heterochromatic AT-rich regions of DNA.

We next tested whether the FTC models could correctly classify live neurons. The models failed to classify live neurons derived from P19 and ES cells (Figure S5G). These results suggest that two different models trained on live cultured cells and fixed tissue cells are necessary for their classification.

### Creating a pan-neuronal classifier

To examine whether the FTC models could sort other subtypes of neurons into the group of neurons, we tested the models on three untrained subtypes of neurons: mitral cells, which are TBR1-positive (TBR1^+^) neurons in the embryonic olfactory bulb (OB), ^33^ embryonic trigeminal ganglion (TG) neurons, which are sensory neurons of the peripheral nervous system, ^34^ and Purkinje cells, which are GABAergic neurons in the postnatal cerebellum (Figures 3A-3D). ^35^ The average accuracies for correctly sorting OB neurons, TG neurons and Purkinje cells were 83.95 ± 4.31%, 2.42 ± 0.85% and 8.79 ± 3.74%, respectively. These accuracies were much lower than that for CTX neurons (Figure 2E).

**Figure 3.**
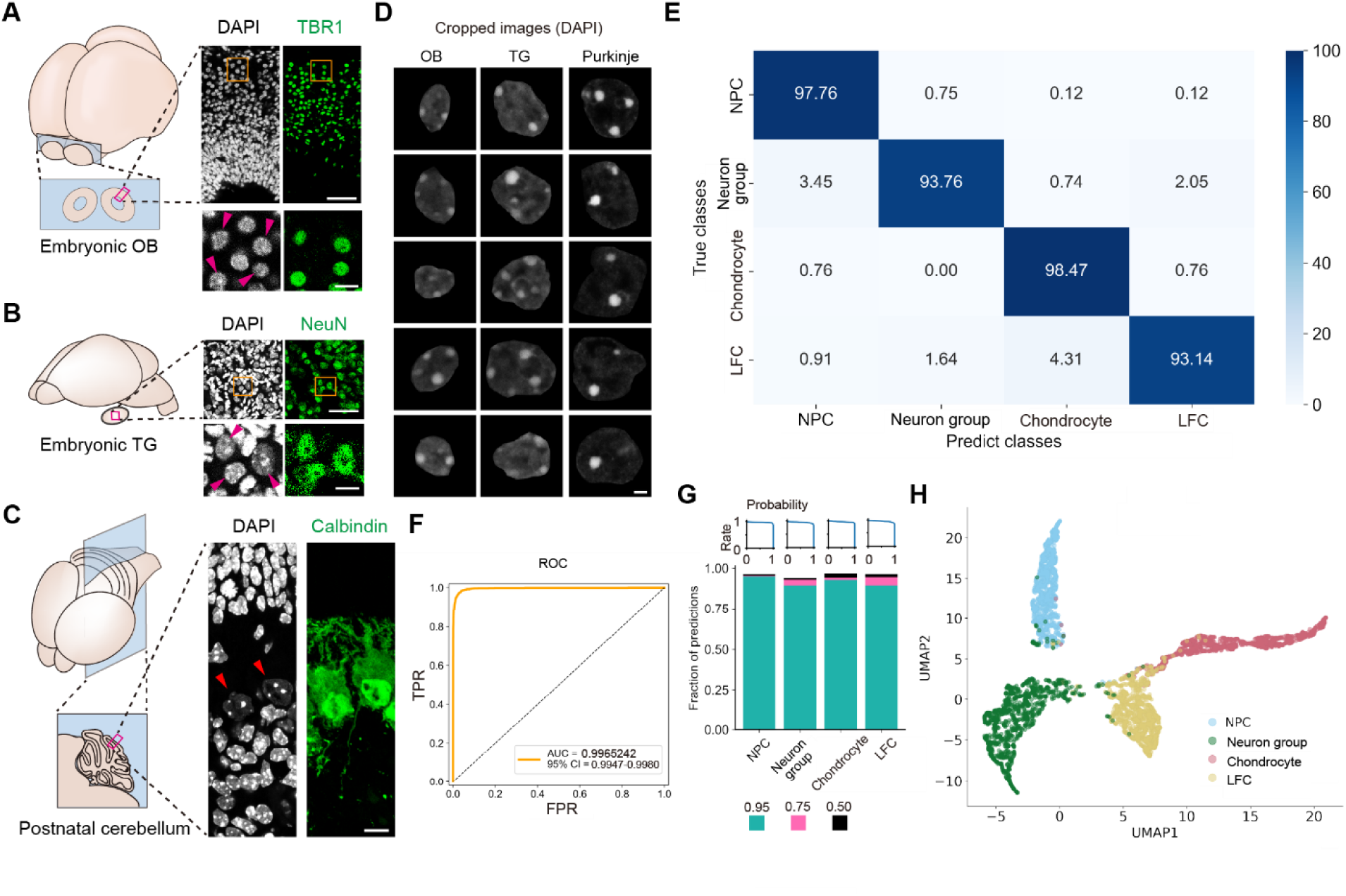
Cell type classification using a unified group of neurons. (A-C) Schematic drawings (left) and representative immunostaining images (right) of E13.5 OB (A), E15.5 TG (B) and postnatal day 4 (P4) cerebellum (C). Magenta boxes are magnified. NeuN and calbindin were used as markers for neurons and Purkinje cells, respectively. Scale bars, 50 µm (top) and 10 µm (bottom) (A and B), 10 µm (C). (D) Representative images of cropped nuclei. Scale bar, 2 µm. (E) Confusion matrix of predictions on an external test set (NPCs, *n =* 201; neuron group, *n =* 916; chondrocytes, *n =* 213; LFCs, *n =* 206). (F) ROC curve and AUC for a representative model (model 5 in Table S3). (G) Top, the fraction (*y* axis) that was correctly classified at or above a certain confidence (probability) threshold (*x* axis) for each cell type. Bottom, the fraction of each cell type that was correctly classified with a confidence score above certain thresholds (>0.95, green; >0.75 and ≤0.95, magenta; >0.50 and ≤0.75, black). (H) UMAP representation of nuclear features learned by the model 5.

We investigated whether a better model could be generated that is able to sort untrained subtypes of neurons into a single unified neuron group, by incorporating those subtypes of neurons into the training set. To maintain data balance, we adjusted the total number of images in this combined neuron dataset (CTX neurons, *n =* 4530; OB neurons, *n =* 4515; TG neurons, *n =* 4515; Purkinje cells, *n =* 4515) to those of other three cell types (NPCs, *n =* 18120; chondrocytes, *n =* 18480; LFCs, *n =* 18120). We used the same CNN architecture with stratified eight-fold cross-validation as above, and the eight models showed high and consistent performance, achieving an average accuracy of 95.80 ± 0.29% on validation sets (Figures S6A and S6B; Table S3) and 94.86 ± 0.79% on an external test set (Figure 3E). Within the unified neuron group, the average accuracies for correctly sorting CTX neurons, OB neurons, TG neurons and Purkinje cells were 93.50 ± 2.55%, 91.98 ± 1.24%, 90.45 ± 3.21% and 98.59 ± 0.80%, respectively. The AUC remained high, at nearly 1.000 (Figure 3F). Most predictions were made with a high confidence score above 0.95 (Figure 3G). UMAP analysis showed four distinct clusters, each corresponding to one of the four cell types (Figure 3H). Zooming into the neuron group cluster revealed overlapping distributions of the four subtypes of neurons (Figure S6C), suggesting that the models may have learned some features common to those subtypes of neurons.

To assess the generalization capability of the models, we tested three untrained subtypes of neurons, TBR1^+^, CDP^+^ and LHX6^+^ neurons in the postnatal CTX, which have distinct functional properties and gene expression profiles (Figures 4A and 4B). ^29,36,37^ Whereas TBR1^+^ and CDP^+^ neurons are glutamatergic, LHX6^+^ neurons are GABAergic. The models sorted the three subtypes of neurons into the unified neuron group with an average accuracy of 95.96 ± 1.36% (Figure 4C). Most predictions were made with a high confidence score above 0.95 (Figure 4D). UMAP analysis showed that the vast majority of all the subtypes of neurons clustered tightly in the neuron group region (Figures 4E and S7). Hereafter the models are designated as unified fixed neuron (UFN) models. These results suggest that the models may be able to sort even untrained subtypes of neurons into the unified neuron group, based on some common nuclear features among subtypes of neurons.

**Figure 4.**
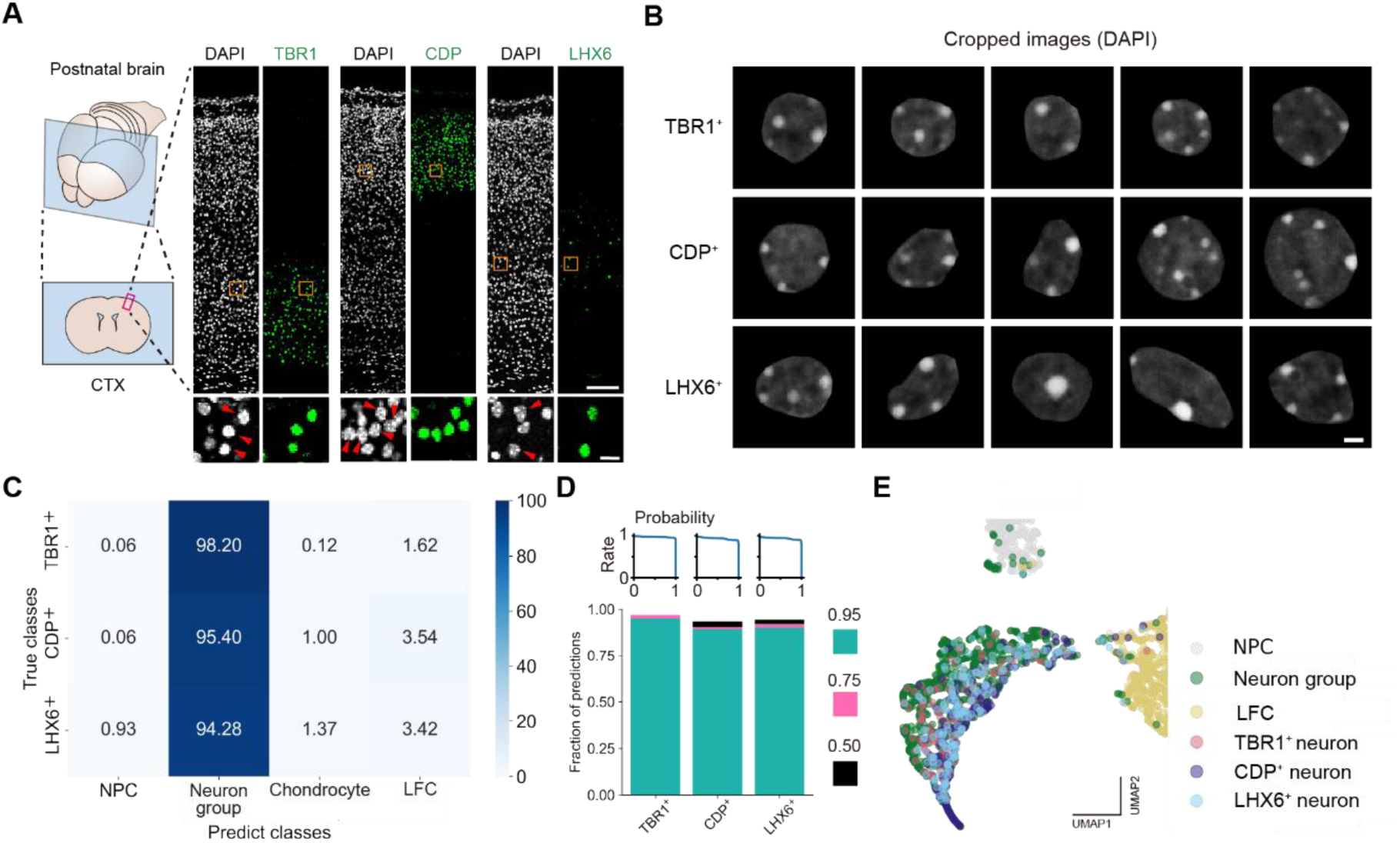
Generalization of the UFN models for untrained subtypes of neurons. (A) P4 CTX: schematic drawing (left) and representative immunostaining images (right). Scale bars, 100 µm (top), 10 µm (bottom). Magenta boxes are magnified. (B) Representative images of cropped nuclei. Scale bar, 2 µm. (C) Average accuracies of the UFN models on TBR1^+^, CDP^+^ and LHX6^+^ neurons (*n =*201 for each subtype of neurons). (D) Top, the fraction (*y* axis) that was correctly categorized at or above a certain confidence (probability) threshold (*x* axis) for each subtype of neurons. Bottom, the fraction of each subtype that was correctly categorized with a confidence score above certain thresholds (>0.95, green; >0.75 and ≤0.95, magenta; >0.50 and ≤0.75, black). (E) Magnified view of the UMAP space around the neuron group region in Figure 3H. The nuclear features of three untrained subtypes of neurons are additionally plotted (*n* = 201 for each subtype of neurons).

### Robustness of cell type classification even in genetic perturbation

We examined the robustness of the UFN models, using genetically manipulated neurons and NPCs by overexpression of key regulatory genes for differentiation with *in vivo* electroporation. ^38^ The transfected cells expressing enhanced yellow fluorescent protein (EYFP) as a control were detected in both the ventricular zone (VZ), where the majority of NPCs reside, and the cortical plate (CP), where neurons migrate to (Figures 5A-5C). In contrast, overexpression of the dominant-negative form of *Maml1* (*dnMaml1*) and constitutively active form of *Notch* (*caNotch*) caused precocious differentiation of neurons and inhibited neuronal differentiation, respectively, as previously described ^39^ EYFP^+^ NPCs and EYFP^+^ neurons were greatly reduced by the overexpression of *dnMaml1* and *caNotch*, respectively. Note that some NPCs and neurons were EYFP^-^, because some cells were not transfected. The UFN models successfully sorted both manipulated neurons and NPCs as well as control ones with an average accuracy of 94.30 ± 1.99% (Figure 5D). Most predictions were made with a high confidence score above 0.95 (Figure 5E). UMAP analysis showed that manipulated neurons and NPCs were segregated in two distinct regions with control ones (Figure 5F). These findings suggest that nuclear features detectable by deep learning are maintained even after the perturbation of neuronal differentiation.

**Figure 5.**
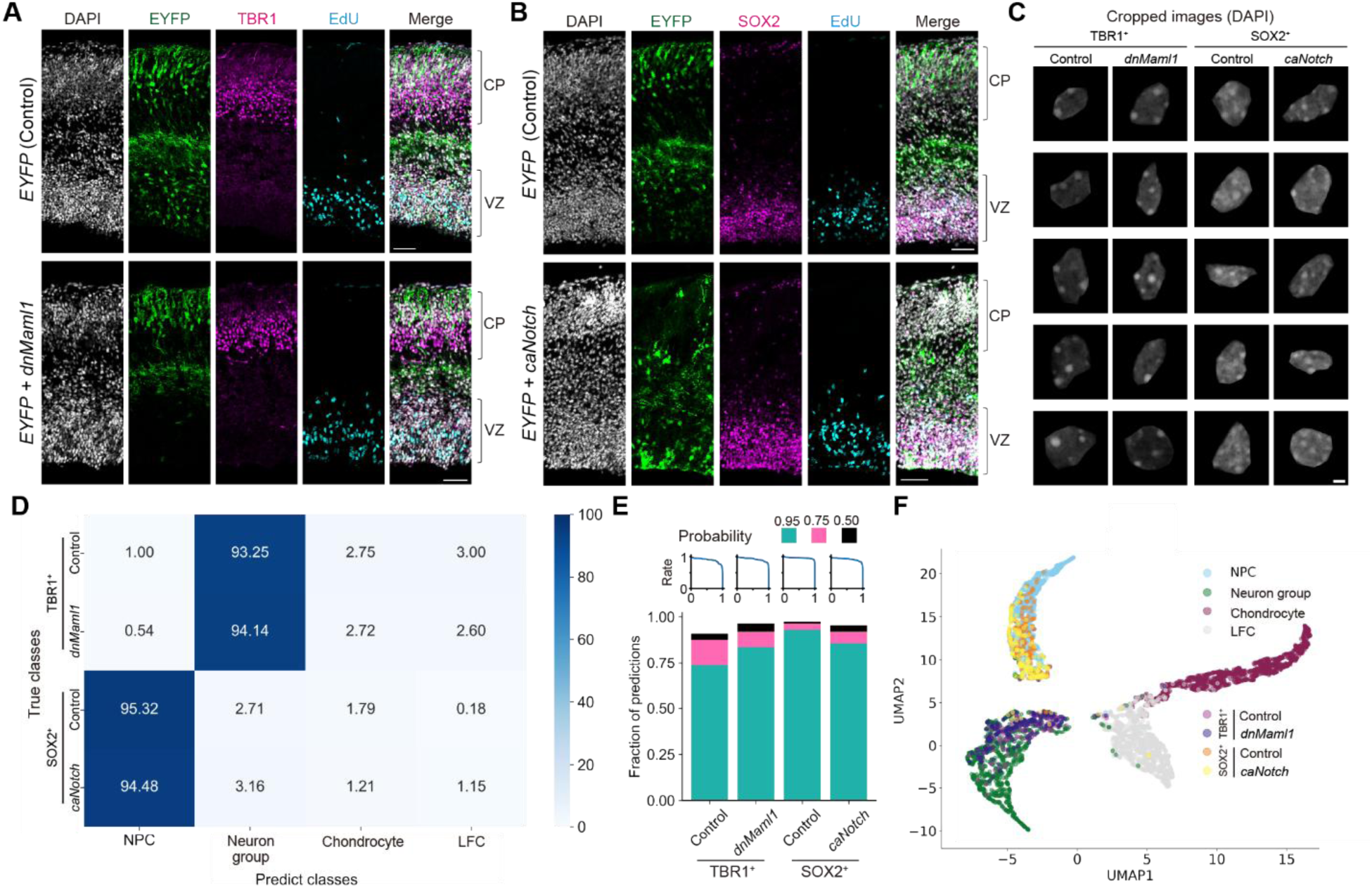
Robustness of the UFN models to genetic perturbation. (A and B) Representative immunostaining images of E16.5 CTX, three days after electroporation of *EYFP* alone (top) and with *dnMaml1* or *caNotch* (bottom). Scale bars, 50 µm. (C) Representative images of cropped nuclei of EYFP^+^TBR1^+^ neurons and EYFP^+^SOX2^+^ NPCs. Scale bar, 2 µm. (D) Average accuracies of the UFN models on transfected cells (EYFP^+^TBR1^+^ neurons in control (*n =* 200) and *dnMaml1* overexpression (*n =* 207), EYFP^+^SOX2^+^ NPCs in control (*n =* 203) and *caNotch* overexpression (*n =* 206)). (E) Top, the fraction (*y* axis) that was correctly classified at or above a certain confidence (probability) threshold (*x* axis) for each cell type. Bottom, the fraction of each cell type that was correctly classified with a confidence score above certain thresholds (>0.95, green; >0.75 and ≤0.95, magenta; >0.50 and ≤0.75, black). (F) The nuclear features of the transfected cells are additionally plotted on Figure 3H.

## DISCUSSION

In this study, we have demonstrated that high-resolution two-dimensional single-plane images of Hoechst- or DAPI-stained nuclei serve as cell-type-specific identifiers for live cultured cells and fixed tissue cells. Our findings suggest that the deep-learning-decipherable distribution patterns of Hoechst and DAPI that bind to DNA are conserved among the same cell type and maintained throughout the nucleus. Despite huge varieties of subtypes of neurons including morphologies and functional properties, unified grouping of various subtypes of neurons by the UFN models might be linked with pan-neuronal features, possibly because neurons share expression of neuron-specific genes for their functions.

Different models were required for live cultured cells and fixed tissue cells. The nuclear organization of fixed cells may have been affected by sample preparation, in which a harsh antigen retrieval step is required to unmask epitopes for immunostaining. ^39^

Defining the precise nuclear patterns that enable cell type classification remains a challenge. To identify cell-type-specific patterns, we applied integrated gradients (IG), ^40^ which visualizes feature attributions by showing important features for each model prediction as a saliency map (Figures S8-S10), but no consistent and interpretable patterns were observed for both live and fixed cells. This may suggest that our CNN models do not classify cell types simply relying on easily discernible features. The optical resolution of our confocal microscope (∼200 nm in a single plane) is insufficient to resolve the positions of individual dye molecules bound to DNA, implying that the CNN models may extract some larger features of nuclear DNA staining. Future efforts focusing on the interpretability of the models will be essential for deciphering the fundamental principles of nuclear organization underlying cell types.

## RESOURCE AVAILABILITY

### Lead contact

For further inquiries, protocol details, or requests for resources, please contact the lead contact, Tetsuichiro Saito.

### Materials availability

This study did not generate new unique reagents.

### Data and Code availability

All image data used for RF and CNN models are available from Zenodo (https://doi.org/10.5281/zenodo.17877652). All codes for RF and CNN models were deposited on GitHub (https://github.com/HiromitsuTanaka/Codes_for_RF_and_CNN_models).

## ACKNOWLEDGMENTS

We thank E. Kawakami for helpful discussion, and K. Ishida and T. Morita for technical assistance. This work was supported by JSPS KAKENHI Grant Numbers JP18K19369 to T.S. and JP24K09655 to H.T.

## AUTHOR CONTRIBUTIONS

Conceptualization and Project administration: T.S.; Investigation, Methodology and Writing & editing: K.S., H.T. and T.S.

## EXPERIMENTAL MODEL AND STUDY PARTICIPANT DETAILS

### Mice

All experimental procedures were approved by the Animal Care and Use Committee of Chiba University and conducted in accordance with the Guidelines for Use of Laboratory Animals (Japan Neuroscience Society) and the ARRIVE guidelines. ICR mice (CLEA Japan) were used for all experiments. The noon of a day when a vaginal plug was detected was designated as E0.5, and the day of birth was designated as P0.

### Cell culture

P19 cells were maintained and differentiated into neurons as previously described with slight modifications. ^22^ Dulbecco’s modified Eagle’s medium (DMEM) (Gibco, 10566016) and 1 µM all-trans retinoic acid (Sigma-Aldrich, R2625) were used. Four days after retinoic acid treatment, cell aggregates were washed in Dulbecco’s phosphate-buffered saline (D-PBS) (Wako, 045-29795), treated with 0.25% trypsin-EDTA solution (Gibco, 25200056) for 3 min at room temperature (RT), dissociated into single cells by gentle pipetting and plated onto glass-base dishes (Iwaki, 3970-035) coated with 0.1% poly(ethyleneimine) (Sigma-Aldrich, P3143) in Neurobasal medium (Gibco, 21103049) containing 1% GlutaMAX (Gibco, 35050061), 2% B-27 supplement (Gibco, 17504044), 10 ng/ml brain-derived neurotrophic factor (Wako, 020-12913) and 1% penicillin/streptomycin (P/S) (Wako, 168-23191). Mouse astrocytes were cultured as previously described with slight modifications. ^41^ Briefly, hippocampi were dissected out from P1 pups, treated with 0.25% trypsin (Gibco, 15090046), dissociated by trituration and cultured on cell culture dishes (Falcon, 353002) in astrocyte medium (Gibco, A1261301) containing 1% P/S. For live-cell analyses, the cells were seeded on glass-base dishes coated with 0.1% poly-D-lysine (Sigma-Aldrich, P7280 and P6407). NIH3T3 fibroblasts ^21^ were seeded on glass-base dishes in DMEM supplemented with 10% fetal bovine serum (FBS) and 1% P/S.

Mouse ES cells (AES0139:EB3) were obtained from RIKEN Bioresource Research Center and cultured as described in the protocols with slight modifications. ^42,43^ Briefly, the cells were maintained in Glasgow’s MEM (Gibco, 11710035) supplemented with 1% FBS, 10% KnockOut Serum Replacement (Gibco, 10828028), 1 mM sodium pyruvate (Gibco, 11360070), 1x Non-Essential Amino Acids (Gibco, 11140050), 0.1 mM 2-mercaptoethanol (Gibco, 21985023), 1000 U/ml recombinant mouse LIF (Sigma-Aldrich, ESG1106) and 0.5% P/S on glass-base dishes coated with 0.1% gelatin (Sigma-Aldrich, ES-006-B). They were differentiated into neurons using the neuronal differentiation medium RHB-A (Takara, Y40001) according to the manufacturer’s protocol. ^44^

## METHOD DETAILS

### Live-cell staining

All live cultured cells were stained with Hoechst (Invitrogen, H3570) and NeuO (NeuroFluor NeuO, Veritas, ST-01801) to detect neurons, as described previously with slight modifications. ^23,45^ Briefly, cells were incubated in the medium containing 125 nM NeuO for 1 h and then the medium containing 5 µg/ml Hoechst for 15 min at 37°C. Then the cells were washed three times with the medium containing no Hoechst or NeuO before live-cell imaging.

### Immunohistochemistry

Tissue preparation and immunostaining were performed as described previously with slight modifications. ^39^ Briefly, animals were weighed and injected with EdU (100 mg/kg, Invitrogen, E10187) in PBS 2 h before tissue collection. All samples were fixed in 4% paraformaldehyde in PBS on ice for the following duration: 2 h for E13.5 heads, 3 h for E15.5 heads, 2.5 h for E16.5 brains, and overnight for P4 brains. Tissues were then cryoprotected in 30% sucrose in PBS overnight, embedded in OCT compound (Sakura, 4583) and stored at −80°C. Cryosections (10—12 µm) were prepared using a cryostat (Leica Biosystems, CM3050S). Sagittal sections were prepared from the cerebellum, whereas coronal sections were prepared from all other tissues.

To unmask epitopes for immunostaining, glass slides containing tissue sections were incubated with TE buffer (10 mM Tris, 1 mM EDTA, pH9.0) for 1 min at 95℃. The slides were washed three times with Tris-buffered saline (TBS), blocked with blocking buffer (0.5% skim milk, 0.25% fish gelatin, 0.5% Triton-X 100 in TBS) for 1 h at RT and incubated with primary antibodies overnight at 4℃ in the blocking buffer. After washing the slides three times with TBS, EdU staining was performed using Click-IT EdU Cell Proliferation Kit (Invitrogen, C10340 or C10637) according to the manufacturer’s protocol. Then the slides were washed once with TBS and four times with washing buffer (1% donkey serum (Sigma-Aldrich, S30-100ML), 0.05% Triton-X 100 in TBS), incubated with secondary antibodies containing 4 µg/ml DAPI (Roche, 10236276001) or 20 µg/ml Hoechst in the blocking buffer for 90 min at RT. After washing twice with the washing buffer and five times with TBS, the slides were covered with VECTASHIELD Vibrance Antifade Mounting Medium (Vector Laboratories, H-1700) and cover glasses (Matsunami glass, No.1S grade).

The following primary antibodies were used: goat anti-SOX2 (1:800, R&D systems, AF2018), goat anti-GFP (1:500, Abcam, ab6673), rabbit anti-TBR1 (1:400, Abcam, ab31940 for embryonic tissues) and (1:200, Sigma-Aldrich, AB10554 for postnatal tissues), rabbit anti-CDP (1:200, Santa Cruz, sc-13024), rabbit anti-Calbindin (1:200, Swant, CB-38a), rabbit anti-GFP (1:500, Invitrogen, A-11122), mouse anti-LHX6 (1:200, Santa Cruz, sc-271433) and mouse anti-NeuN (1:400, Sigma-Aldrich, MAB377).

The following secondary antibodies were used: donkey anti-goat IgG conjugated with DyLight 488 (Abcam, ab96935), donkey anti-goat IgG conjugated with DyLight 550 (Abcam, ab96936), donkey anti-goat IgG conjugated with DyLight 650 (Abcam, ab96938), donkey anti-rabbit IgG conjugated with DyLight 488 (Abcam, ab96919), donkey anti-rabbit IgG conjugated with DyLight 550 (Abcam, ab96920) and donkey anti-mouse IgG conjugated with DyLight 550 (Abcam, ab98795). The dilution ratio for all secondary antibodies was 1:250.

### Alcian Blue staining

Chondrocytes were stained with Alcian Blue (Sigma-Aldrich, A5268) as described previously with slight modifications. ^30^ Briefly, glass slides containing tissue sections were incubated in 3% acetic acid for 3 min and in 0.1% Alcian Blue, 3% acetic acid for 5 min at RT. Then the slides were washed once with 3% acetic acid, five times with distilled water and once with TBS.

### *In vivo* electroporation

We used plasmids pCAG-EYFP ^38^ as a control to express EYFP, and pCAG-EYFP-CAG-caNotch ^38^ and pCAG-EYFP-CAG-dnMaml1 ^39^ to express *caNotch* and *dnMaml1*, respectively, with EYFP. *In vivo* electroporation was performed as described previously with slight modification. ^38,46^ Briefly, pregnant mice at E13.5 were anesthetized with intraperitoneal injection with medetomidine (0.75 mg/kg, Nippon Zenyaku Kogyo), midazolam (4 mg/kg, Maruishi Seiyaku) and butorphanol tartrate (5 mg/kg, Meiji Animal Health) in saline. After injection of the plasmids, electric pulses were delivered with forceps-type electrodes. The electroporated brains were dissected out at E16.5.

### Fluorescent microscopy

Fluorescent images of live or fixed cells were taken using a confocal laser-scanning microscope (Leica, TCS SP8) equipped with a 20× objective lens, a 63× oil-immersion objective lens, a 160× oil-immersion objective lens (Leica, Optik Kit GSD, numerical aperture 1.43), Hybrid detectors and 405-, 488-, 552- and 638-nm laser lines. We used optimized instrument settings to exclude spectral crosstalk between different fluorophores. We acquired 8-bit grayscale images (4096 × 4096 pixels) using the 160× objective lens to build RF and CNN models. Live-cell imaging with the SP8 confocal microscope was performed using a stage-top incubator (Tokai Hit, STXG-GSI2X-SET) to maintain conditions at 37°C and 5% CO₂.

Phase contrast and fluorescent images of live cells were acquired using a confocal laser-scanning microscope (Olympus, FV10i-LIV) equipped with a 10× objective lens or an inverted microscope (Olympus, CKX41) equipped with a 40× objective lens.

### Nuclear data

We manually cropped one to six images from each nucleus using ImageJ (National Institutes of Health). When multiple images were cropped from the same nucleus, we chose images the focal planes of which were separated by at least 0.56 µm. The cropped images were centered on a black background large enough to cover the whole nucleus, with a size of 1248 × 1248 pixels for live-cell images or 720 × 720 pixels for fixed-cell images. For external test sets, we cropped images from nuclei that were not used for training.

### RF classification

For building RF models, we quantified twelve nuclear morphological features using ImageJ: Area, Perimeter, Width, Height, Major, Minor, Circularity, Solidity, Aspect ratio, Feret, MinFeret and Feret angle. We built RF models using RandomForestClassifier from the Scikit-learn Python package and performed stratified *k*-fold cross-validation ^47,48^. The nuclear images were resized to 150 × 150 pixels before feature extraction. The parameters of the RF model (max features, n_estimator and *k*-fold) were tuned to improve the prediction performance. The optimal parameter values were determined to be max features = 0.8, n_estimator = 300 and *k* = 5.

### CNN model architecture and training

We constructed CNN models using multiple convolution layers, max pooling layers and dropout layers (Figure S3A). After flattening, FC layers were employed. Each of convolution and FC layers, except for the output layer, was followed by the rectified linear unit activation function. The output layer utilized the softmax activation function. We adopted batch normalization twice. Model parameters were optimized using the Adam optimizer with a learning rate of 1×10^-3^ and decay of 1×10^-3^. Categorical cross-entropy was used as the loss function. Models were trained for 100 epochs.

We generated additional images by adjusting the brightness of each image within a range of 0.6 to 1.0, with increments of 0.1. Furthermore, we augmented the dataset using ImageDataGenerator from Keras. Each image was augmented by applying random rotations within a range of 180 degrees and random zooms within a range of 0.7 to 1.3. Horizontal and vertical flips were also applied. All input images, following augmentation, were resized to 150 × 150 pixels. We performed stratified eight-fold cross-validation. ^48^

### Model interpretability and visualization

To visualize the high-dimensional features learned by the CNN models, we applied UMAP using the UMAP-learn Python library. The 128-dimensional output of the final FC layer was projected onto a two-dimensional plane, with each point representing a single nucleus. To generate saliency maps for model interpretation, we applied IG using custom Python scripts adapted from the original implementation (https://github.com/PAIR-code/saliency). ^40^

### Statistical analysis

For multiclass classification, the predicted class was defined as the class with the highest predicted probability score. To evaluate classification performance, we calculated the accuracy, precision, recall, F1-score and AUC for each class using one-vs-rest approach based on the results from stratified *k*-fold cross-validation. ^48^ Further, we computed micro average, macro average and weighted average for each metric using R (v4.4.1, The R Project for Statistical Computing). Nonparametric bootstrapping with 2,000 samples was used to compute 95% confidence intervals (CI). Statistical significance of pairwise comparison was detected by Welch’s ANOVA with post-hoc Games-Howell’s test using Prism8 (GraphPad Software). All data in the text are presented as mean ± standard deviation (s.d.).

