## Supplemental Figures and Tables for "Deep learning cell type classification using nuclear DNA patterns"

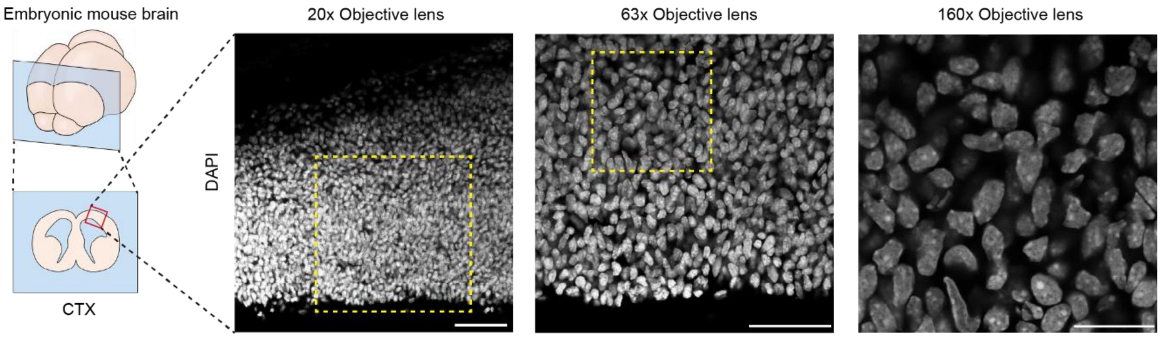


**Figure S1. Nuclear images at higher magnifications. Related to Figure 1.**

Embryonic day 13.5 (E13.5) mouse brain coronal section: schematic drawing (left), DAPI-stained cerebral cortex (CTX) section at different magnifications (right). The yellow dashed boxes indicate the regions imaged at higher magnifications. Scale bars, 50 µm (20× and 63× Objective lenses), 20 µm (160× Objective lens).

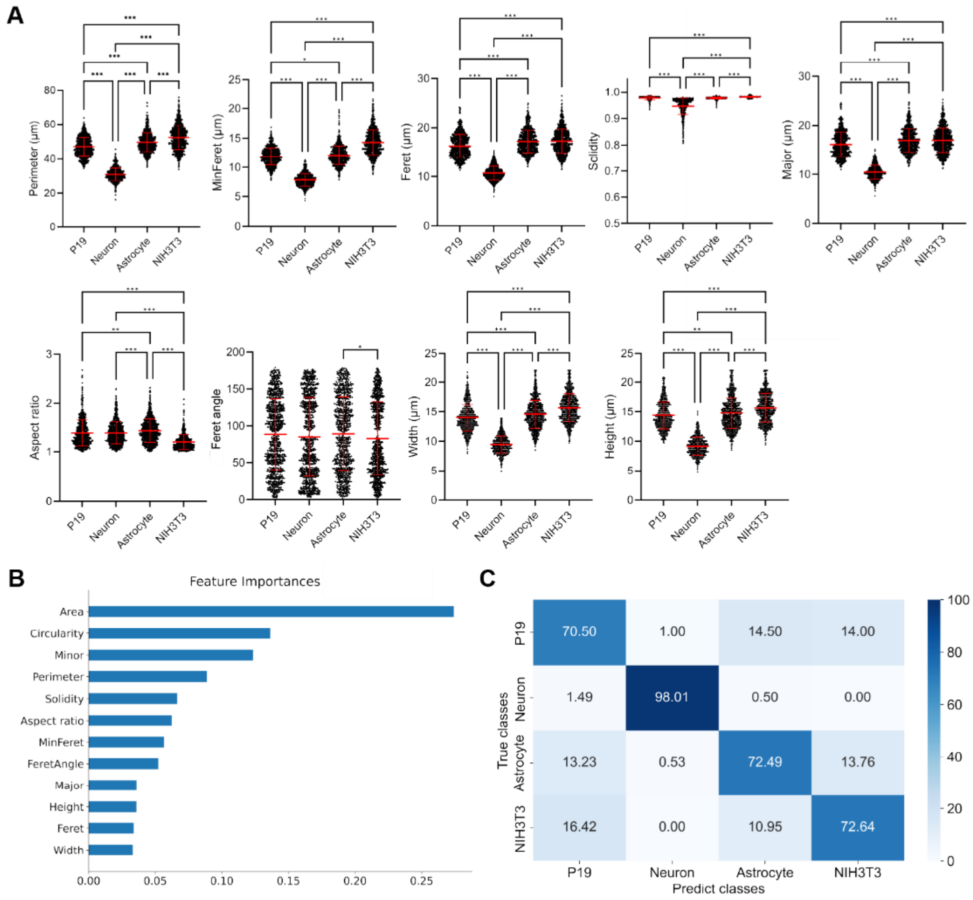


**Figure S2. RF model analysis of nuclear morphology. Related to Figure 1.**

(A) Quantification of nine nuclear morphological features (see Figure 1C for the other features) for each cell type. Mean and s.d. are shown. Statistical significance was assessed by Welch's ANOVA with post-hoc Games-Howell's test. *$p < 0.05$, **$p < 0.01$, ***$p < 0.001$.

(B) Feature importance scores from a representative model trained using five-fold cross-validation.

(C) Confusion matrix for the RF model on the validation set.

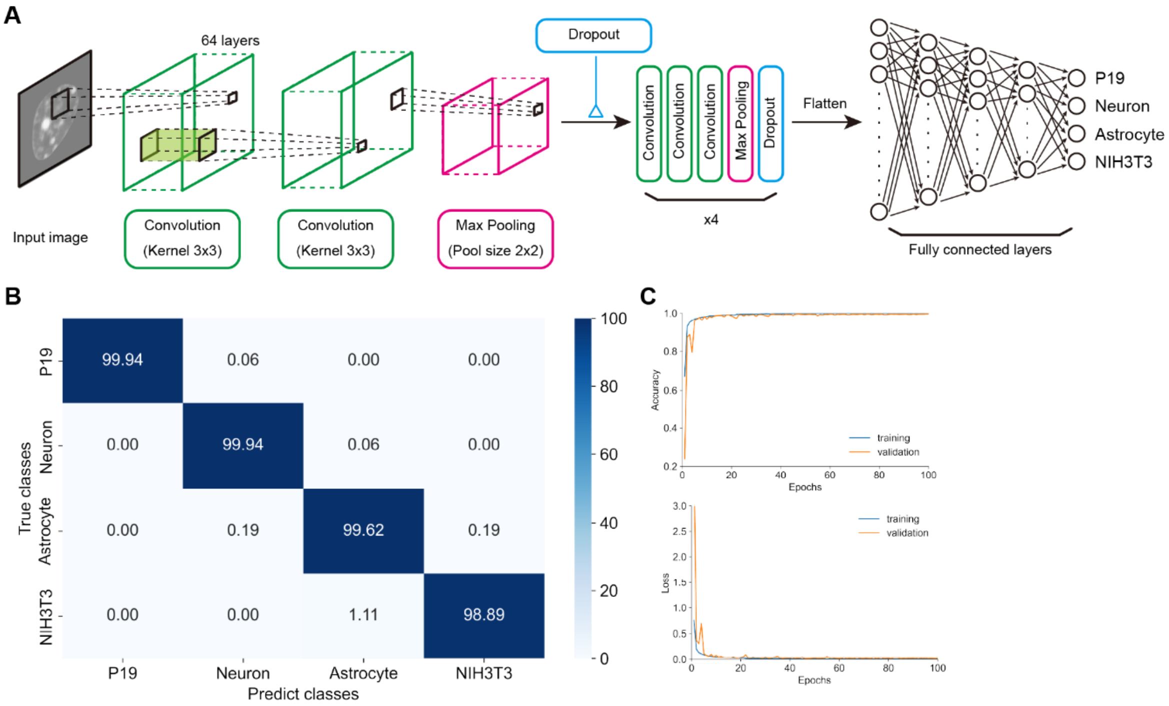


**Figure S3. CNN architecture and model performance on live cultured cells. Related to Figure 1.**

(A) Schematic of the CNN architecture. The CNN models were trained on nuclear images of live cells (150 × 150 pixels, pixel size: 0.148 µm).

(B) Confusion matrix for a representative model (model 6 in Table S1) on the validation set.

(C) Learning curves through the CNN training. Accuracy (top) and loss (bottom) in the training set and the validation set for the model 6 over 100 epochs.

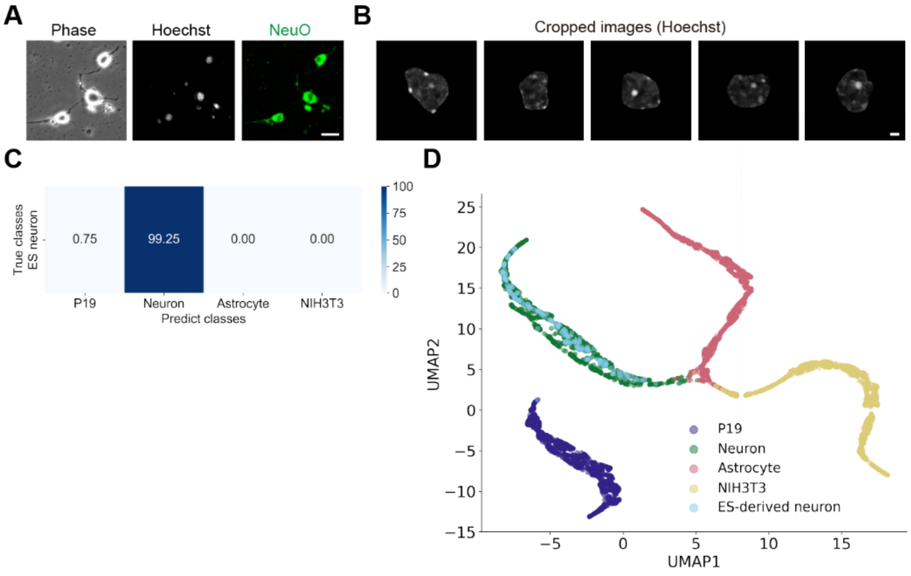


**Figure S4. ES cell-derived neuron classification. Related to Figure 1.**

(A) Representative phase-contrast and fluorescent (Hoechst and NeuO staining) images of ES cell-derived neurons. Scale bar, 20 µm.

(B) Representative nuclei images. Scale bar: 2 µm.

(C) Average accuracy of the LCC models on ES cell-derived neurons (n = 200).

(D) The nuclear features of the ES cell-derived neurons are additionally plotted on Figure 1G (n = 200).

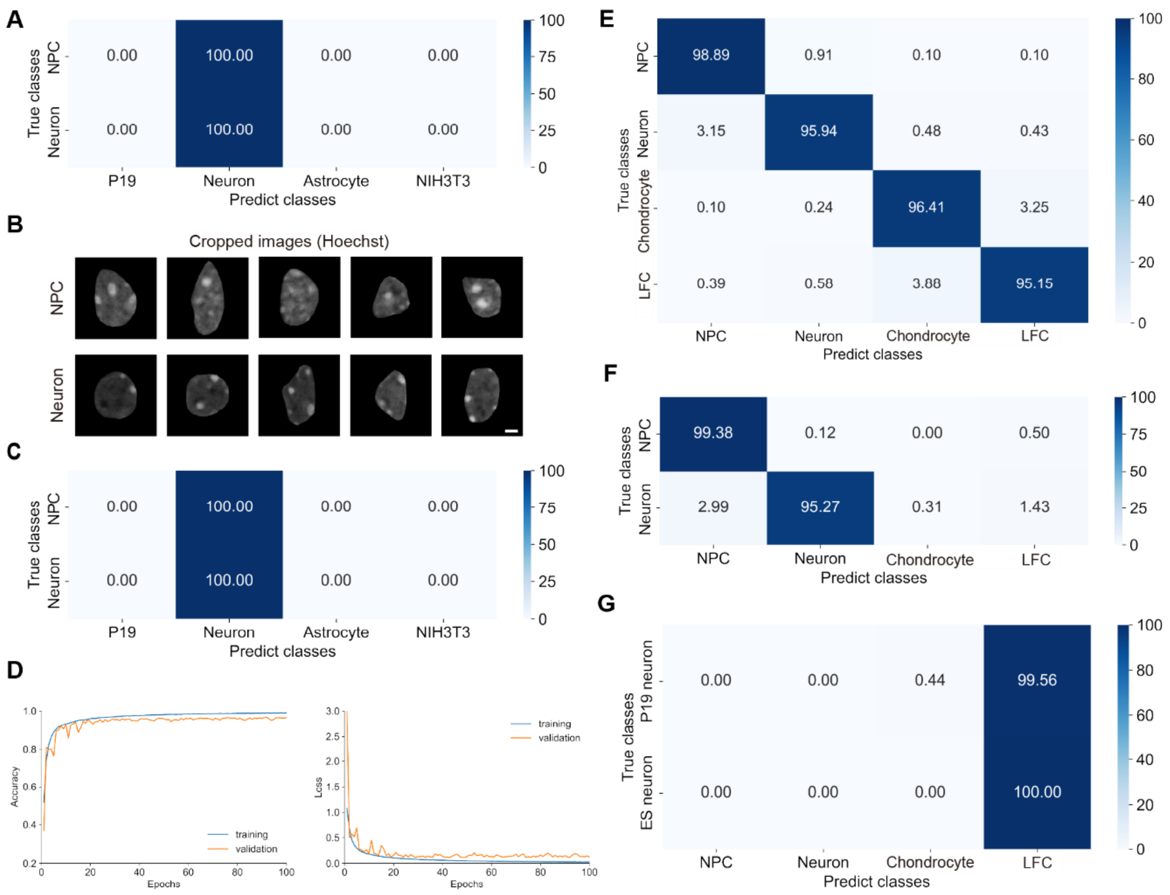


**Figure S5. The LCC and FTC models are not interchangeable. Related to Figure 2.**

(A and C) Average accuracies of the LCC models on fixed NPCs and neurons stained with DAPI, which were used in Figure 2E (A) and those stained with Hoechst (n = 201 for each cell type) (C).

(B) Representative nuclei images of the Hoechst-stained nuclei. Scale bar, 2 µm.

(D) Learning curves through the CNN training on cells in fixed tissues. Accuracy (left) and loss (right) in the training set and the validation set for a representative model (model 7 in Table S2) over 100 epochs.

(E) Confusion matrix for the model 7 on the validation set.

(F and G) Average accuracies of the FTC models on fixed NPCs and neurons stained with Hoechst, which were used in Figure S5C (F) and live neurons derived from P19 and ES cells, which were used in Figures 1E and S4C (G). For the test set, cropped nuclei were centered on a 1248 × 1248 pixels (A, C) or 720 × 720 pixels (F, G) black background and resized to 150 × 150 pixels to match the input format of the LCC and FTC models, respectively.

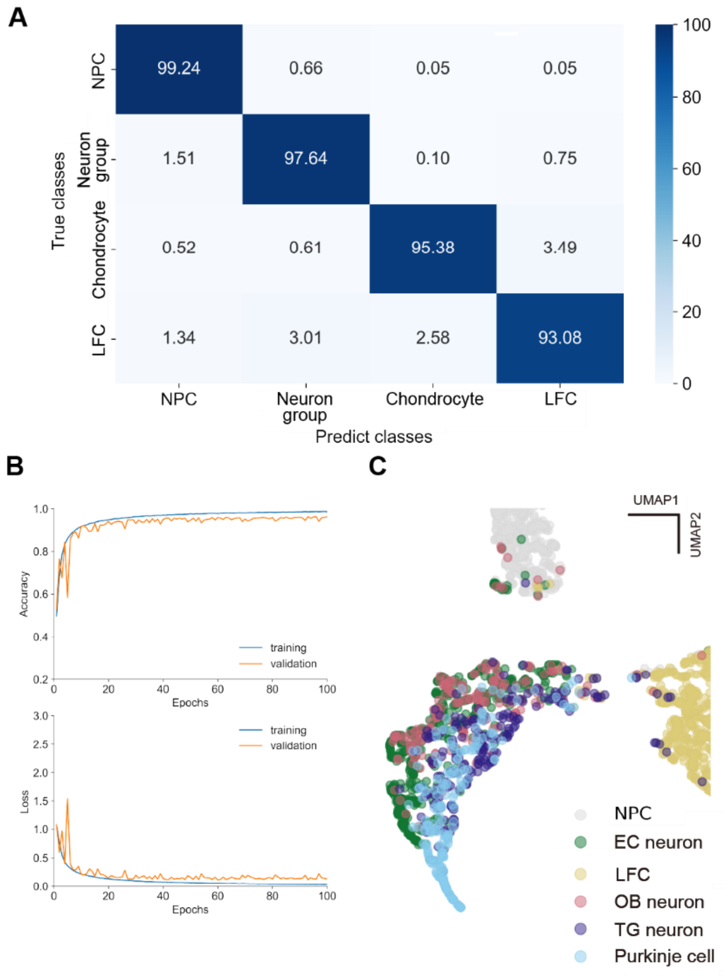


**Figure S6. Model performance on a unified neuron group. Related to Figure 3.**

(A) Confusion matrix for a representative model (model 5 in Table S3) on the validation set.

(B) Learning curves through the CNN training. Accuracy (top) and loss (bottom) in the training set and the validation set for the model 5 over 100 epochs.

(C) Magnified view of the UMAP space around the neuron group region in Figure 3H. The nuclear features of the four subtypes of neurons are plotted (Embryonic CTX (EC) neurons, n = 522; OB neurons, n = 215; TG neurons, n = 212; Purkinje cells, n = 239).

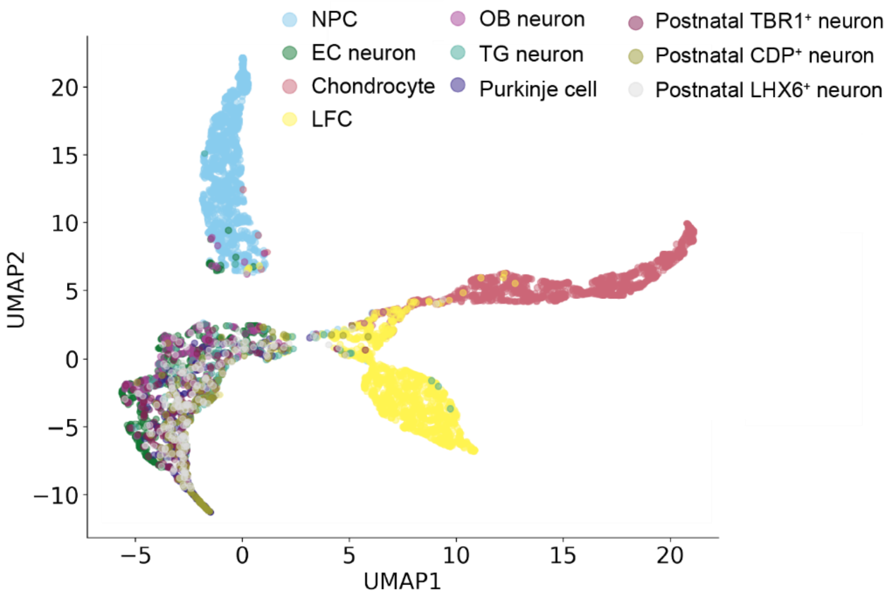


**Figure S7. UMAP representation of seven subtypes of neurons. Related to Figure 4.**

The nuclear features of three more subtypes of neurons are plotted on Figure 3H.

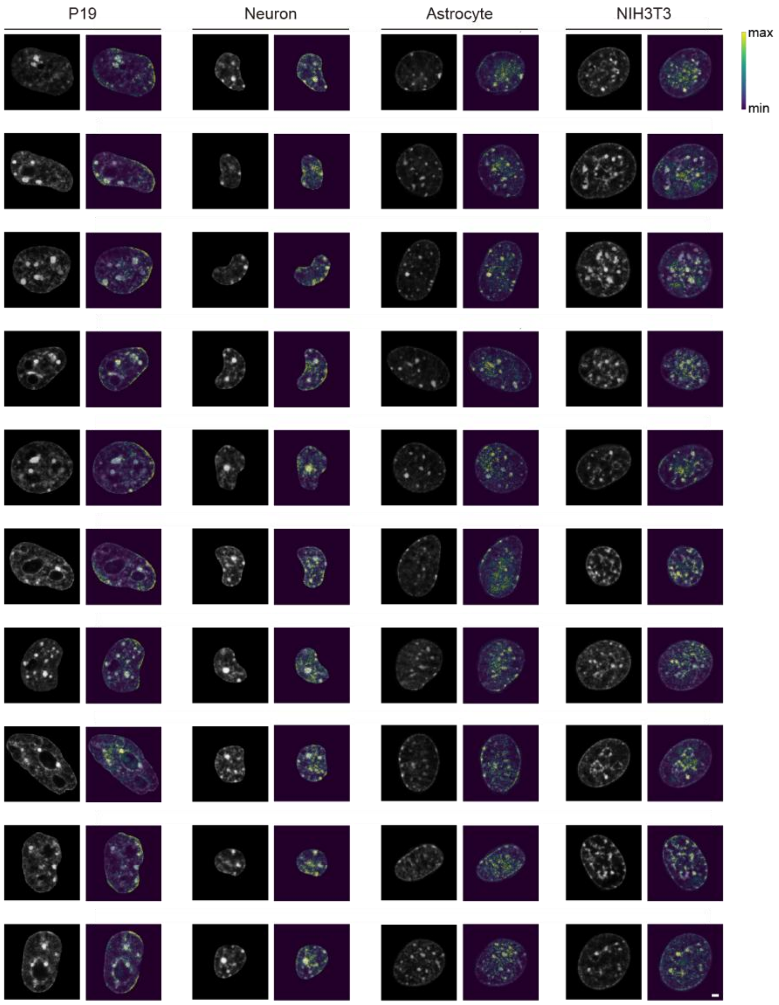


**Figure S8. Visualization of IGs for live-cell classification using the LCC model. Related to Figure 1.**

Ten examples per cell type showing input nuclear images with (right) and without (left) the visualizations obtained from IGs for the model 6 in Table S1. Those cells were correctly classified. Yellow and blue regions in IGs indicate positive and minimal contributions to prediction, respectively. Scale bar, 2 µm.

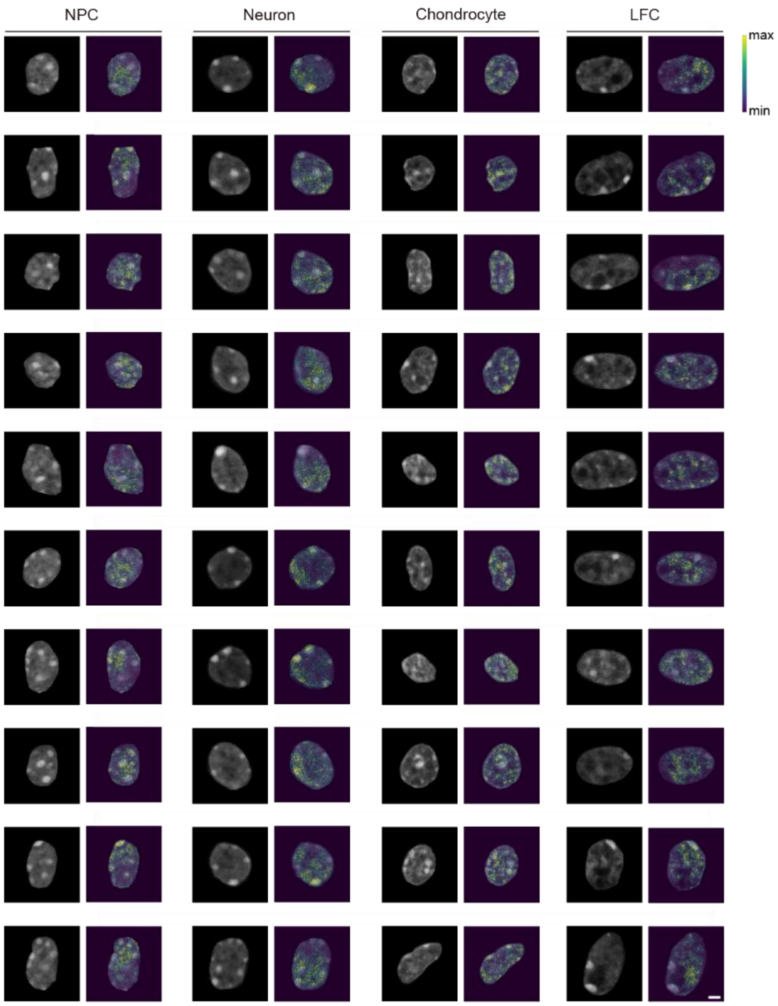


**Figure S9. Visualization of IGs for fixed-cell classification using the FTC model. Related to Figure 2.**

Ten examples per cell type showing input nuclear images with (right) and without (left) the visualizations obtained from IGs for the model 7 in Table S2. Those cells were correctly classified. Yellow and blue regions in IGs indicate positive and minimal contributions to prediction, respectively. Scale bar, 2 µm.

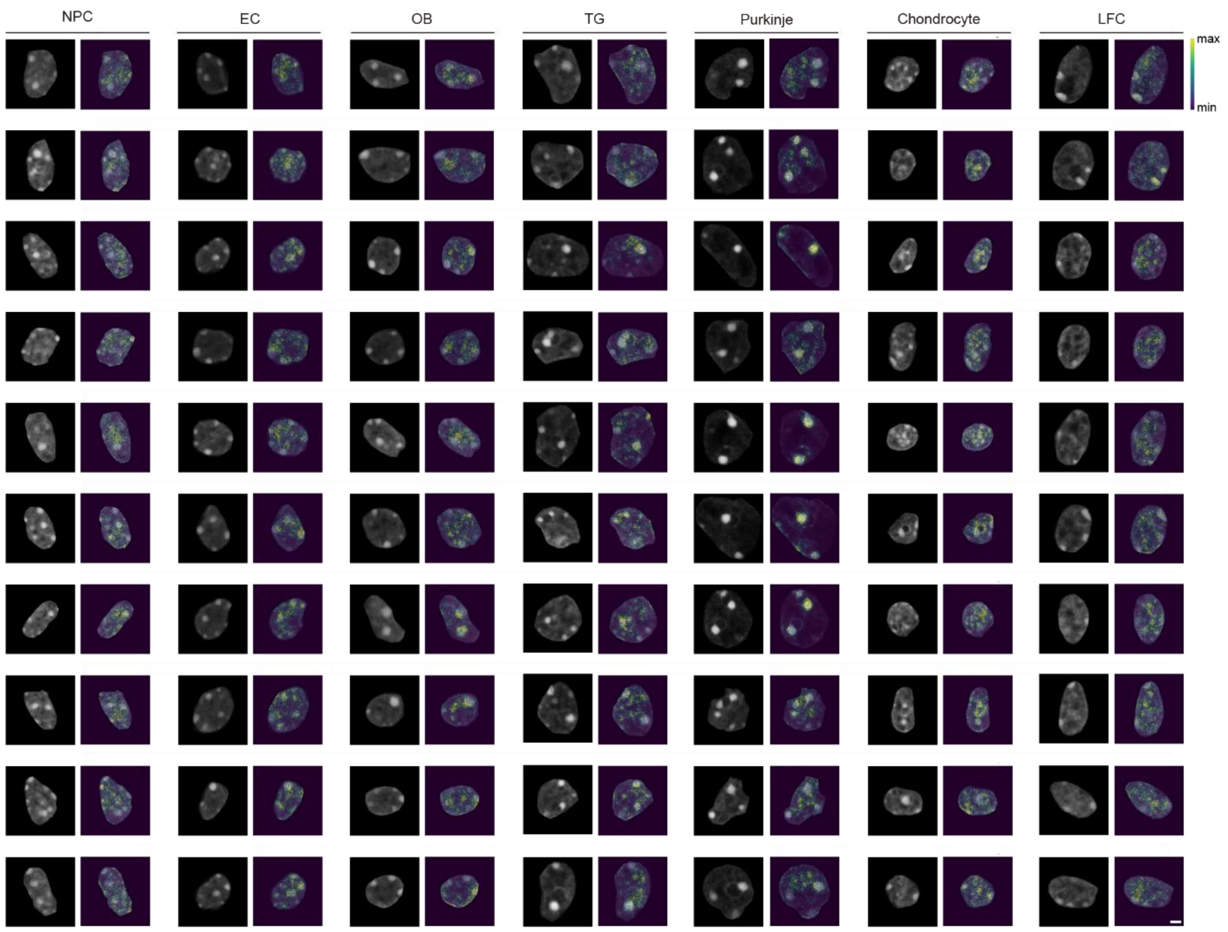


**Figure S10. Visualization of IGs for fixed-cell classification using the UFN model. Related to Figure 3.**

Ten examples per cell type and per subtype of neurons showing input nuclear images with (right) and without (left) the visualizations obtained from IGs for the model 5 in Table S3. Those cells were correctly classified. Yellow and blue regions in IGs indicate positive and minimal contributions to prediction, respectively. Scale bar, 2 µm.

| | model 1 | | | | model 2 | | | |
|---|---|---|---|---|---|---|---|---|
| | Precision | Recall | F1-score | AUC | Precision | Recall | F1-score | AUC |
| P19 | 1.0000 | 0.9970 | 0.9985 | 0.9999 | 0.9970 | 0.9988 | 0.9979 | 1.0000 |
| | 1.0000-1.0000 | 0.9943-0.9994 | 0.9971-0.9997 | 0.9998-1.0000 | 0.9941-0.9994 | 0.9970-1.0000 | 0.9964-0.9994 | 1.0000-1.0000 |
| Neuron | 0.9825 | 0.9964 | 0.9894 | 0.9995 | 0.9976 | 0.9893 | 0.9935 | 0.9999 |
| | 0.9760-0.9885 | 0.9934-0.9989 | 0.9857-0.9929 | 0.9990-0.9999 | 0.9952-0.9994 | 0.9841-0.9939 | 0.9906-0.9960 | 0.9998-1.0000 |
| Astrocyte | 0.9949 | 0.9687 | 0.9816 | 0.9994 | 0.9821 | 0.9937 | 0.9879 | 0.9998 |
| | 0.9911-0.9981 | 0.9598-0.9772 | 0.9767-0.9865 | 0.9990-0.9997 | 0.9753-0.9885 | 0.9898-0.9974 | 0.9840-0.9917 | 0.9996-0.9999 |
| NIH3T3 | 0.9826 | 0.9959 | 0.9892 | 0.9999 | 0.9941 | 0.9894 | 0.9917 | 0.9999 |
| | 0.9762-0.9886 | 0.9925-0.9988 | 0.9856-0.9925 | 0.9998-0.9999 | 0.9902-0.9976 | 0.9842-0.9941 | 0.9884-0.9947 | 0.9999-1.0000 |
| Micro avg | 0.9898 | 0.9898 | 0.9898 | 0.9996 | 0.9928 | 0.9928 | 0.9928 | 0.9999 |
| | 0.9873-0.9922 | 0.9873-0.9922 | 0.9873-0.9922 | 0.9993-0.9998 | 0.9907-0.9949 | 0.9907-0.9949 | 0.9907-0.9949 | 0.9998-1.0000 |
| Macro avg | 0.9900 | 0.9895 | 0.9897 | 0.9997 | 0.9927 | 0.9928 | 0.9928 | 0.9999 |
| | 0.9875-0.9923 | 0.9870-0.9919 | 0.9871-0.9921 | 0.9995-0.9999 | 0.9906-0.9948 | 0.9908-0.9949 | 0.9907-0.9948 | 0.9998-0.9999 |
| Weighted avg | 0.9899 | 0.9898 | 0.9898 | 0.9997 | 0.9929 | 0.9928 | 0.9928 | 0.9999 |
| | 0.9875-0.9923 | 0.9873-0.9922 | 0.9873-0.9922 | 0.9995-0.9999 | 0.9908-0.9949 | 0.9907-0.9949 | 0.9907-0.9949 | 0.9998-1.0000 |

| | model 3 | | | | model 4 | | | |
|---|---|---|---|---|---|---|---|---|
| | Precision | Recall | F1-score | AUC | Precision | Recall | F1-score | AUC |
| P19 | 0.9988 | 0.9994 | 0.9991 | 1.0000 | 0.9976 | 1.0000 | 0.9988 | 1.0000 |
| | 0.9969-1.0000 | 0.9982-1.0000 | 0.9979-1.0000 | 1.0000-1.0000 | 0.9952-0.9994 | 1.0000-1.0000 | 0.9976-0.9997 | 1.0000-1.0000 |
| Neuron | 0.9994 | 0.9905 | 0.9949 | 1.0000 | 0.9982 | 0.9917 | 0.9950 | 1.0000 |
| | 0.9981-1.0000 | 0.9859-0.9948 | 0.9925-0.9971 | 0.9999-1.0000 | 0.9958-1.0000 | 0.9870-0.9958 | 0.9924-0.9971 | 0.9999-1.0000 |
| Astrocyte | 0.9770 | 0.9856 | 0.9813 | 0.9996 | 0.9912 | 0.9894 | 0.9903 | 0.9998 |
| | 0.9695-0.9844 | 0.9792-0.9913 | 0.9764-0.9861 | 0.9994-0.9997 | 0.9864-0.9955 | 0.9841-0.9939 | 0.9868-0.9934 | 0.9996-0.9999 |
| NIH3T3 | 0.9865 | 0.9865 | 0.9865 | 0.9997 | 0.9906 | 0.9965 | 0.9935 | 0.9999 |
| | 0.9808-0.9918 | 0.9805-0.9918 | 0.9825-0.9904 | 0.9996-0.9999 | 0.9858-0.9948 | 0.9935-0.9988 | 0.9906-0.9961 | 0.9998-1.0000 |
| Micro avg | 0.9906 | 0.9906 | 0.9906 | 0.9999 | 0.9945 | 0.9945 | 0.9945 | 0.9999 |
| | 0.9880-0.9930 | 0.9880-0.9930 | 0.9880-0.9930 | 0.9998-0.9999 | 0.9927-0.9961 | 0.9927-0.9961 | 0.9927-0.9961 | 0.9999-1.0000 |
| Macro avg | 0.9904 | 0.9905 | 0.9905 | 0.9998 | 0.9944 | 0.9944 | 0.9944 | 0.9999 |
| | 0.9880-0.9928 | 0.9880-0.9929 | 0.9880-0.9929 | 0.9997-0.9999 | 0.9926-0.9961 | 0.9925-0.9961 | 0.9925-0.9961 | 0.9998-1.0000 |
| Weighted avg | 0.9906 | 0.9906 | 0.9906 | 0.9998 | 0.9945 | 0.9945 | 0.9945 | 0.9999 |
| | 0.9881-0.9930 | 0.9880-0.9930 | 0.9880-0.9930 | 0.9997-0.9999 | 0.9927-0.9961 | 0.9927-0.9961 | 0.9927-0.9961 | 0.9999-1.0000 |

| | model 5 | | | | model 6 | | | |
|---|---|---|---|---|---|---|---|---|
| | Precision | Recall | F1-score | AUC | Precision | Recall | F1-score | AUC |
| P19 | 0.9988 | 0.9988 | 0.9988 | 1.0000 | 0.9965 | 0.9988 | 0.9976 | 1.0000 |
| | 0.9969-1.0000 | 0.9970-1.0000 | 0.9974-0.9997 | 1.0000-1.0000 | 0.9934-0.9988 | 0.9970-1.0000 | 0.9958-0.9991 | 1.0000-1.0000 |
| Neuron | 0.9964 | 0.9947 | 0.9956 | 1.0000 | 0.9953 | 0.9947 | 0.9950 | 1.0000 |
| | 0.9933-0.9988 | 0.9909-0.9977 | 0.9932-0.9976 | 0.9999-1.0000 | 0.9916-0.9982 | 0.9907-0.9982 | 0.9924-0.9973 | 1.0000-1.0000 |
| Astrocyte | 0.9857 | 0.9944 | 0.9900 | 0.9999 | 0.9918 | 0.9831 | 0.9874 | 0.9998 |
| | 0.9796-0.9914 | 0.9905-0.9976 | 0.9864-0.9934 | 0.9998-0.9999 | 0.9870-0.9962 | 0.9768-0.9891 | 0.9833-0.9911 | 0.9997-0.9999 |
| NIH3T3 | 0.9959 | 0.9894 | 0.9926 | 0.9999 | 0.9877 | 0.9941 | 0.9909 | 0.9999 |
| | 0.9924-0.9988 | 0.9841-0.9941 | 0.9896-0.9955 | 0.9999-1.0000 | 0.9822-0.9926 | 0.9902-0.9976 | 0.9875-0.9939 | 0.9999-1.0000 |
| Micro avg | 0.9943 | 0.9943 | 0.9943 | 0.9999 | 0.9928 | 0.9928 | 0.9928 | 0.9999 |
| | 0.9924-0.9961 | 0.9924-0.9961 | 0.9924-0.9961 | 0.9999-1.0000 | 0.9907-0.9948 | 0.9907-0.9948 | 0.9907-0.9948 | 0.9999-1.0000 |
| Macro avg | 0.9942 | 0.9943 | 0.9943 | 0.9999 | 0.9928 | 0.9927 | 0.9927 | 0.9999 |
| | 0.9923-0.9960 | 0.9924-0.9961 | 0.9923-0.9960 | 0.9999-1.0000 | 0.9907-0.9948 | 0.9906-0.9946 | 0.9906-0.9947 | 0.9999-1.0000 |
| Weighted avg | 0.9943 | 0.9943 | 0.9943 | 0.9999 | 0.9928 | 0.9928 | 0.9928 | 0.9999 |
| | 0.9924-0.9961 | 0.9924-0.9961 | 0.9924-0.9961 | 0.9999-1.0000 | 0.9907-0.9948 | 0.9907-0.9948 | 0.9907-0.9948 | 0.9999-1.0000 |

| | model 7 | | | | model 8 | | | |
|---|---|---|---|---|---|---|---|---|
| | Precision | Recall | F1-score | AUC | Precision | Recall | F1-score | AUC |
| P19 | 1.0000 | 0.9964 | 0.9982 | 1.0000 | 0.9988 | 0.9959 | 0.9973 | 1.0000 |
| | 1.0000-1.0000 | 0.9933-0.9988 | 0.9966-0.9994 | 0.9999-1.0000 | 0.9970-1.0000 | 0.9926-0.9988 | 0.9955-0.9988 | 0.9999-1.0000 |
| Neuron | 0.9976 | 0.9941 | 0.9958 | 1.0000 | 0.9947 | 0.9941 | 0.9944 | 0.9999 |
| | 0.9951-0.9994 | 0.9901-0.9976 | 0.9935-0.9979 | 0.9999-1.0000 | 0.9911-0.9977 | 0.9902-0.9976 | 0.9916-0.9968 | 0.9997-1.0000 |
| Astrocyte | 0.9912 | 0.9887 | 0.9900 | 0.9998 | 0.9523 | 0.9994 | 0.9753 | 0.9996 |
| | 0.9864-0.9956 | 0.9837-0.9937 | 0.9863-0.9934 | 0.9997-0.9999 | 0.9417-0.9616 | 0.9981-1.0000 | 0.9695-0.9802 | 0.9991-0.9999 |
| NIH3T3 | 0.9878 | 0.9971 | 0.9924 | 0.9999 | 1.0000 | 0.9570 | 0.9780 | 0.9997 |
| | 0.9825-0.9925 | 0.9942-0.9994 | 0.9894-0.9952 | 0.9997-0.9999 | 1.0000-1.0000 | 0.9468-0.9662 | 0.9727-0.9828 | 0.9994-0.9999 |
| Micro avg | 0.9942 | 0.9942 | 0.9942 | 0.9999 | 0.9864 | 0.9864 | 0.9864 | 0.9995 |
| | 0.9924-0.9960 | 0.9924-0.9960 | 0.9924-0.9960 | 0.9999-1.0000 | 0.9835-0.9891 | 0.9835-0.9891 | 0.9835-0.9891 | 0.9992-0.9997 |
| Macro avg | 0.9941 | 0.9941 | 0.9941 | 0.9999 | 0.9864 | 0.9866 | 0.9863 | 0.9998 |
| | 0.9924-0.9959 | 0.9923-0.9959 | 0.9923-0.9959 | 0.9998-1.0000 | 0.9835-0.9890 | 0.9837-0.9893 | 0.9833-0.9889 | 0.9996-0.9999 |
| Weighted avg | 0.9942 | 0.9942 | 0.9942 | 0.9999 | 0.9869 | 0.9864 | 0.9864 | 0.9998 |
| | 0.9924-0.9960 | 0.9924-0.9960 | 0.9924-0.9960 | 0.9998-1.0000 | 0.9843-0.9894 | 0.9835-0.9891 | 0.9835-0.9891 | 0.9996-0.9999 |

**Table S1. Performance metrics of live cultured cell models. Related to Figure 1.**

The one-vs-rest precision, recall, F1-score and AUC for each cell type and the overall metrics (micro, macro and weighted averages) are shown. The 95% CI is indicated below each value.

| | model 1 | | | | model 2 | | | |
|---|---|---|---|---|---|---|---|---|
| | Precision | Recall | F1-score | AUC | Precision | Recall | F1-score | AUC |
| NPC | 0.9872 | 0.9441 | 0.9652 | 0.9987 | 0.9870 | 0.9652 | 0.9759 | 0.9994 |
| | 0.9816-0.9920 | 0.9336-0.9537 | 0.9592-0.9706 | 0.9981-0.9991 | 0.9818-0.9916 | 0.9572-0.9727 | 0.9713-0.9805 | 0.9991-0.9996 |
| Neuron | 0.9555 | 0.9785 | 0.9669 | 0.9986 | 0.9802 | 0.9711 | 0.9756 | 0.9991 |
| | 0.9466-0.9640 | 0.9717-0.9846 | 0.9611-0.9725 | 0.9983-0.9990 | 0.9737-0.9860 | 0.9640-0.9777 | 0.9708-0.9801 | 0.9987-0.9994 |
| Chondrocyte | 0.9148 | 0.9817 | 0.9471 | 0.9965 | 0.9216 | 0.9779 | 0.9489 | 0.9971 |
| | 0.9036-0.9260 | 0.9761-0.9872 | 0.9401-0.9536 | 0.9956-0.9973 | 0.9103-0.9328 | 0.9713-0.9838 | 0.9419-0.9554 | 0.9964-0.9977 |
| LFC | 0.9683 | 0.9151 | 0.9410 | 0.9953 | 0.9605 | 0.9308 | 0.9454 | 0.9956 |
| | 0.9609-0.9758 | 0.9032-0.9267 | 0.9334-0.9484 | 0.9943-0.9962 | 0.9515-0.9688 | 0.9195-0.9416 | 0.9382-0.9523 | 0.9945-0.9966 |
| Micro avg | 0.9550 | 0.9550 | 0.9550 | 0.9970 | 0.9613 | 0.9613 | 0.9613 | 0.9978 |
| | 0.9507-0.9594 | 0.9507-0.9594 | 0.9507-0.9594 | 0.9965-0.9975 | 0.9572-0.9652 | 0.9572-0.9652 | 0.9572-0.9652 | 0.9974-0.9982 |
| Macro avg | 0.9565 | 0.9548 | 0.9550 | 0.9973 | 0.9623 | 0.9612 | 0.9615 | 0.9978 |
| | 0.9523-0.9607 | 0.9504-0.9591 | 0.9507-0.9594 | 0.9968-0.9978 | 0.9583-0.9661 | 0.9570-0.9652 | 0.9574-0.9653 | 0.9973-0.9982 |
| Weighted avg | 0.9563 | 0.9550 | 0.9550 | 0.9973 | 0.9621 | 0.9613 | 0.9614 | 0.9978 |
| | 0.9522-0.9604 | 0.9507-0.9594 | 0.9507-0.9594 | 0.9968-0.9978 | 0.9581-0.9659 | 0.9572-0.9652 | 0.9573-0.9653 | 0.9973-0.9982 |
| | model 3 | | | | model 4 | | | |
| | Precision | Recall | F1-score | AUC | Precision | Recall | F1-score | AUC |
| NPC | 0.9899 | 0.9637 | 0.9766 | 0.9989 | 0.9895 | 0.9667 | 0.9779 | 0.9994 |
| | 0.985-0.994 | 0.9555-0.9715 | 0.9718-0.9812 | 0.9984-0.9994 | 0.9847-0.9939 | 0.9582-0.9738 | 0.9731-0.9822 | 0.9991-0.9996 |
| Neuron | 0.9787 | 0.9687 | 0.9737 | 0.9989 | 0.9771 | 0.9838 | 0.9805 | 0.9969 |
| | 0.9718-0.9847 | 0.9605-0.9761 | 0.9686-0.9785 | 0.9984-0.9992 | 0.9703-0.9834 | 0.9781-0.9890 | 0.9759-0.9847 | 0.9961-0.9977 |
| Chondrocyte | 0.9131 | 0.9909 | 0.9504 | 0.9980 | 0.9331 | 0.9865 | 0.9591 | 0.9978 |
| | 0.9016-0.9246 | 0.9866-0.9947 | 0.9439-0.9571 | 0.9975-0.9985 | 0.9220-0.9433 | 0.9814-0.9916 | 0.9526-0.9649 | 0.9972-0.9984 |
| LFC | 0.9788 | 0.9298 | 0.9537 | 0.9970 | 0.9739 | 0.9328 | 0.9529 | 0.9994 |
| | 0.9719-0.9853 | 0.9182-0.9407 | 0.9469-0.9602 | 0.9962-0.9978 | 0.9664-0.9810 | 0.9221-0.9436 | 0.9457-0.9593 | 0.9992-0.9996 |
| Micro avg | 0.9634 | 0.9634 | 0.9634 | 0.9979 | 0.9676 | 0.9676 | 0.9676 | 0.9984 |
| | 0.9593-0.9674 | 0.9593-0.9674 | 0.9593-0.9674 | 0.9974-0.9983 | 0.9635-0.9713 | 0.9635-0.9713 | 0.9635-0.9713 | 0.9980-0.9987 |
| Macro avg | 0.9651 | 0.9633 | 0.9636 | 0.9982 | 0.9684 | 0.9675 | 0.9676 | 0.9984 |
| | 0.9612-0.9689 | 0.9592-0.9672 | 0.9595-0.9676 | 0.9978-0.9986 | 0.9645-0.9721 | 0.9634-0.9712 | 0.9635-0.9714 | 0.9980-0.9987 |
| Weighted avg | 0.9649 | 0.9634 | 0.9635 | 0.9982 | 0.9682 | 0.9676 | 0.9676 | 0.9984 |
| | 0.9611-0.9686 | 0.9593-0.9674 | 0.9594-0.9676 | 0.9978-0.9986 | 0.9643-0.9719 | 0.9635-0.9713 | 0.9635-0.9713 | 0.9980-0.9987 |
| | model 5 | | | | model 6 | | | |
| | Precision | Recall | F1-score | AUC | Precision | Recall | F1-score | AUC |
| NPC | 0.9932 | 0.9382 | 0.9649 | 0.9983 | 0.9904 | 0.9666 | 0.9784 | 0.9993 |
| | 0.9893-0.9969 | 0.9274-0.9490 | 0.9589-0.9709 | 0.9976-0.9989 | 0.9859-0.9945 | 0.9589-0.9747 | 0.974-0.983 | 0.9990-0.9996 |
| Neuron | 0.9469 | 0.9868 | 0.9664 | 0.9955 | 0.9800 | 0.9843 | 0.9822 | 0.9995 |
| | 0.9369-0.9569 | 0.9815-0.9914 | 0.9606-0.9721 | 0.9944-0.9965 | 0.9739-0.9862 | 0.9787-0.9895 | 0.9781-0.9863 | 0.9993-0.9996 |
| Chondrocyte | 0.9205 | 0.9803 | 0.9495 | 0.9969 | 0.9370 | 0.9808 | 0.9584 | 0.9980 |
| | 0.9087-0.9315 | 0.9738-0.9860 | 0.9426-0.9560 | 0.9962-0.9975 | 0.9265-0.9467 | 0.9746-0.9864 | 0.9518-0.9643 | 0.9975-0.9985 |
| LFC | 0.9736 | 0.9220 | 0.9471 | 0.9986 | 0.9667 | 0.9397 | 0.9530 | 0.9973 |
| | 0.9663-0.9806 | 0.9103-0.9329 | 0.9398-0.9537 | 0.9982-0.9990 | 0.9587-0.9740 | 0.9292-0.9494 | 0.9463-0.9595 | 0.9967-0.9979 |
| Micro avg | 0.9569 | 0.9569 | 0.9569 | 0.9967 | 0.9679 | 0.9679 | 0.9679 | 0.9985 |
| | 0.9524-0.9613 | 0.9524-0.9613 | 0.9524-0.9613 | 0.9961-0.9972 | 0.9641-0.9717 | 0.9641-0.9717 | 0.9641-0.9717 | 0.9982-0.9988 |
| Macro avg | 0.9586 | 0.9568 | 0.9570 | 0.9973 | 0.9685 | 0.9678 | 0.9680 | 0.9985 |
| | 0.9541-0.9628 | 0.9523-0.9610 | 0.9525-0.9613 | 0.9968-0.9978 | 0.9648-0.9722 | 0.9641-0.9717 | 0.9642-0.9718 | 0.9982-0.9988 |
| Weighted avg | 0.9584 | 0.9569 | 0.9569 | 0.9973 | 0.9684 | 0.9679 | 0.9679 | 0.9985 |
| | 0.9542-0.9625 | 0.9524-0.9613 | 0.9524-0.9613 | 0.9968-0.9978 | 0.9647-0.9721 | 0.9641-0.9717 | 0.9642-0.9717 | 0.9982-0.9988 |
| | model 7 | | | | model 8 | | | |
| | Precision | Recall | F1-score | AUC | Precision | Recall | F1-score | AUC |
| NPC | 0.9889 | 0.9627 | 0.9756 | 0.9990 | 0.9874 | 0.9598 | 0.9734 | 0.9992 |
| | 0.9842-0.9933 | 0.9540-0.9704 | 0.9707-0.9802 | 0.9986-0.9994 | 0.9824-0.9921 | 0.9516-0.9676 | 0.9683-0.9780 | 0.9990-0.9995 |
| Neuron | 0.9594 | 0.9829 | 0.9710 | 0.9989 | 0.9681 | 0.9814 | 0.9747 | 0.9992 |
| | 0.9505-0.9672 | 0.9772-0.9884 | 0.9655-0.9760 | 0.9986-0.9993 | 0.9602-0.9759 | 0.9754-0.9870 | 0.9698-0.9794 | 0.9989-0.9994 |
| Chondrocyte | 0.9641 | 0.9557 | 0.9599 | 0.9974 | 0.9213 | 0.9798 | 0.9497 | 0.9969 |
| | 0.9561-0.9716 | 0.9466-0.9640 | 0.9538-0.9655 | 0.9967-0.9981 | 0.9100-0.9321 | 0.9735-0.9858 | 0.9428-0.9563 | 0.9961-0.9976 |
| LFC | 0.9515 | 0.9617 | 0.9566 | 0.9972 | 0.9700 | 0.9210 | 0.9449 | 0.9961 |
| | 0.9418-0.9603 | 0.9534-0.9697 | 0.9499-0.9625 | 0.9964-0.9979 | 0.9627-0.9772 | 0.9090-0.9325 | 0.9374-0.9520 | 0.995-0.997 |
| Micro avg | 0.9657 | 0.9657 | 0.9657 | 0.9981 | 0.9606 | 0.9606 | 0.9606 | 0.9978 |
| | 0.9616-0.9694 | 0.9616-0.9694 | 0.9616-0.9694 | 0.9977-0.9985 | 0.9564-0.9647 | 0.9564-0.9647 | 0.9564-0.9647 | 0.9973-0.9982 |
| Macro avg | 0.9660 | 0.9658 | 0.9658 | 0.9981 | 0.9617 | 0.9605 | 0.9607 | 0.9979 |
| | 0.9618-0.9696 | 0.9616-0.9694 | 0.9616-0.9694 | 0.9977-0.9985 | 0.9577-0.9656 | 0.9562-0.9646 | 0.9564-0.9648 | 0.9974-0.9983 |
| Weighted avg | 0.9659 | 0.9657 | 0.9657 | 0.9981 | 0.9615 | 0.9606 | 0.9606 | 0.9978 |
| | 0.9618-0.9696 | 0.9616-0.9694 | 0.9616-0.9694 | 0.9977-0.9985 | 0.9575-0.9655 | 0.9564-0.9647 | 0.9564-0.9648 | 0.9974-0.9983 |

**Table S2. Performance metrics of fixed tissue cell models. Related to Figure 2.**

The one-vs-rest precision, recall, F1-score and AUC for each cell type and the overall metrics (micro, macro and weighted averages) are shown. The 95% CI is indicated below each value.

| | model 1 | | | | model 2 | | | |
|---|---|---|---|---|---|---|---|---|
| | Precision | Recall | F1-score | AUC | Precision | Recall | F1-score | AUC |
| NPC | 0.9876 | 0.9735 | 0.9805 | 0.9996 | 0.9829 | 0.9863 | 0.9846 | 0.9996 |
| | 0.9826-0.9920 | 0.9660-0.9806 | 0.9761-0.9847 | 0.9994-0.9997 | 0.9771-0.9881 | 0.9812-0.9913 | 0.9808-0.9884 | 0.9994-0.9997 |
| Neuron group | 0.9730 | 0.9549 | 0.9638 | 0.9976 | 0.9772 | 0.9460 | 0.9614 | 0.9972 |
| | 0.9659-0.9798 | 0.9460-0.9641 | 0.9578-0.9698 | 0.9969-0.9983 | 0.9704-0.9834 | 0.9362-0.9554 | 0.9554-0.9674 | 0.9961-0.9982 |
| Chondrocyte | 0.9561 | 0.9534 | 0.9547 | 0.9974 | 0.9184 | 0.9851 | 0.9506 | 0.9973 |
| | 0.9470-0.9648 | 0.9441-0.9621 | 0.9479-0.9611 | 0.9968-0.9980 | 0.9071-0.9295 | 0.9796-0.9899 | 0.9439-0.9572 | 0.9966-0.9980 |
| LFC | 0.9162 | 0.9490 | 0.9323 | 0.9941 | 0.9502 | 0.9068 | 0.9280 | 0.9931 |
| | 0.9038-0.9278 | 0.9392-0.9583 | 0.9241-0.9400 | 0.9930-0.9952 | 0.9400-0.9589 | 0.8943-0.9189 | 0.9197-0.9359 | 0.9916-0.9944 |
| Micro avg | 0.9577 | 0.9577 | 0.9577 | 0.9974 | 0.9562 | 0.9562 | 0.9562 | 0.9968 |
| | 0.9533-0.9619 | 0.9533-0.9619 | 0.9533-0.9619 | 0.9970-0.9979 | 0.9518-0.9606 | 0.9518-0.9606 | 0.9518-0.9606 | 0.9961-0.9974 |
| Macro avg | 0.9582 | 0.9577 | 0.9579 | 0.9972 | 0.9572 | 0.9561 | 0.9561 | 0.9968 |
| | 0.9540-0.9625 | 0.9533-0.9620 | 0.9535-0.9621 | 0.9967-0.9976 | 0.9530-0.9615 | 0.9517-0.9605 | 0.9518-0.9606 | 0.9962-0.9974 |
| Weighted avg | 0.9582 | 0.9577 | 0.9578 | 0.9972 | 0.9570 | 0.9562 | 0.9561 | 0.9968 |
| | 0.9540-0.9624 | 0.9533-0.9619 | 0.9535-0.9621 | 0.9967-0.9976 | 0.9528-0.9613 | 0.9518-0.9606 | 0.9517-0.9605 | 0.9961-0.9974 |

| | model 3 | | | | model 4 | | | |
|---|---|---|---|---|---|---|---|---|
| | Precision | Recall | F1-score | AUC | Precision | Recall | F1-score | AUC |
| NPC | 0.9881 | 0.9745 | 0.9813 | 0.9992 | 0.9791 | 0.9858 | 0.9824 | 0.9993 |
| | 0.9832-0.9926 | 0.9676-0.9810 | 0.9769-0.9852 | 0.9987-0.9995 | 0.9723-0.9850 | 0.9802-0.9906 | 0.9780-0.9863 | 0.9987-0.9997 |
| Neuron group | 0.9723 | 0.9474 | 0.9597 | 0.9970 | 0.9527 | 0.9700 | 0.9613 | 0.9955 |
| | 0.9647-0.9793 | 0.9378-0.9571 | 0.9532-0.9659 | 0.9963-0.9977 | 0.9436-0.9615 | 0.9621-0.9767 | 0.9552-0.9671 | 0.9945-0.9964 |
| Chondrocyte | 0.9270 | 0.9760 | 0.9508 | 0.9976 | 0.9422 | 0.9716 | 0.9567 | 0.9977 |
| | 0.9154-0.9379 | 0.9691-0.9821 | 0.9442-0.9572 | 0.9970-0.9981 | 0.9324-0.9511 | 0.9646-0.9785 | 0.9505-0.9627 | 0.9971-0.9982 |
| LFC | 0.9328 | 0.9191 | 0.9259 | 0.9930 | 0.9642 | 0.9095 | 0.9361 | 0.9978 |
| | 0.9217-0.9431 | 0.9074-0.9305 | 0.9173-0.9341 | 0.9917-0.9942 | 0.9557-0.9722 | 0.8966-0.9215 | 0.9278-0.9438 | 0.9971-0.9984 |
| Micro avg | 0.9544 | 0.9544 | 0.9544 | 0.9968 | 0.9593 | 0.9593 | 0.9593 | 0.9977 |
| | 0.9497-0.9589 | 0.9497-0.9589 | 0.9497-0.9589 | 0.9963-0.9973 | 0.9550-0.9635 | 0.9550-0.9635 | 0.9550-0.9635 | 0.9973-0.9981 |
| Macro avg | 0.9550 | 0.9543 | 0.9544 | 0.9967 | 0.9595 | 0.9592 | 0.9591 | 0.9976 |
| | 0.9506-0.9593 | 0.9498-0.9586 | 0.9499-0.9588 | 0.9961-0.9972 | 0.9553-0.9637 | 0.9550-0.9635 | 0.9548-0.9634 | 0.9971-0.9980 |
| Weighted avg | 0.9549 | 0.9544 | 0.9544 | 0.9967 | 0.9595 | 0.9593 | 0.9591 | 0.9976 |
| | 0.9505-0.9592 | 0.9497-0.9589 | 0.9498-0.9589 | 0.9961-0.9972 | 0.9552-0.9637 | 0.9550-0.9635 | 0.9548-0.9634 | 0.9971-0.9980 |

| | model 5 | | | | model 6 | | | |
|---|---|---|---|---|---|---|---|---|
| | Precision | Recall | F1-score | AUC | Precision | Recall | F1-score | AUC |
| NPC | 0.9925 | 0.9662 | 0.9792 | 0.9994 | 0.9920 | 0.9696 | 0.9807 | 0.9994 |
| | 0.9883-0.9960 | 0.9584-0.9739 | 0.9746-0.9835 | 0.9992-0.9996 | 0.9877-0.9956 | 0.9618-0.9769 | 0.9761-0.9849 | 0.9990-0.9997 |
| Neuron group | 0.9764 | 0.9562 | 0.9662 | 0.9952 | 0.9626 | 0.9612 | 0.9619 | 0.9980 |
| | 0.9700-0.9826 | 0.9469-0.9647 | 0.9603-0.9717 | 0.9942-0.9961 | 0.9543-0.9707 | 0.9530-0.9696 | 0.9561-0.9679 | 0.9973-0.9986 |
| Chondrocyte | 0.9534 | 0.9726 | 0.9629 | 0.9975 | 0.9175 | 0.9832 | 0.9492 | 0.9976 |
| | 0.9443-0.9621 | 0.9655-0.9792 | 0.9570-0.9686 | 0.9969-0.9981 | 0.9056-0.9283 | 0.9773-0.9884 | 0.9423-0.9557 | 0.9970-0.9981 |
| LFC | 0.9308 | 0.9554 | 0.9429 | 0.9981 | 0.9505 | 0.9040 | 0.9267 | 0.9941 |
| | 0.9191-0.9411 | 0.9461-0.9637 | 0.9350-0.9496 | 0.9976-0.9987 | 0.9411-0.9599 | 0.8909-0.9157 | 0.9180-0.9342 | 0.9929-0.9951 |
| Micro avg | 0.9627 | 0.9627 | 0.9627 | 0.9977 | 0.9546 | 0.9546 | 0.9546 | 0.9973 |
| | 0.9584-0.9667 | 0.9584-0.9667 | 0.9584-0.9667 | 0.9972-0.9981 | 0.9502-0.9590 | 0.9502-0.9590 | 0.9502-0.9590 | 0.9967-0.9977 |
| Macro avg | 0.9632 | 0.9626 | 0.9628 | 0.9976 | 0.9557 | 0.9545 | 0.9546 | 0.9973 |
| | 0.9591-0.9672 | 0.9584-0.9667 | 0.9585-0.9668 | 0.9971-0.9980 | 0.9513-0.9598 | 0.9502-0.9588 | 0.9502-0.9589 | 0.9967-0.9977 |
| Weighted avg | 0.9632 | 0.9627 | 0.9628 | 0.9976 | 0.9555 | 0.9546 | 0.9546 | 0.9973 |
| | 0.9590-0.9672 | 0.9584-0.9667 | 0.9586-0.9668 | 0.9971-0.9980 | 0.9513-0.9597 | 0.9502-0.9590 | 0.9501-0.9590 | 0.9967-0.9977 |

| | model 7 | | | | model 8 | | | |
|---|---|---|---|---|---|---|---|---|
| | Precision | Recall | F1-score | AUC | Precision | Recall | F1-score | AUC |
| NPC | 0.9876 | 0.9794 | 0.9835 | 0.9996 | 0.9944 | 0.9598 | 0.9768 | 0.9991 |
| | 0.9827-0.9923 | 0.9729-0.9852 | 0.9795-0.9872 | 0.9994-0.9997 | 0.9909-0.9975 | 0.9511-0.9682 | 0.9719-0.9813 | 0.9986-0.9995 |
| Neuron group | 0.9729 | 0.9523 | 0.9625 | 0.9976 | 0.9508 | 0.9690 | 0.9598 | 0.9977 |
| | 0.9655-0.9799 | 0.9430-0.9610 | 0.9563-0.9683 | 0.9968-0.9983 | 0.9414-0.9600 | 0.9613-0.9768 | 0.9539-0.9660 | 0.9971-0.9983 |
| Chondrocyte | 0.9424 | 0.9750 | 0.9584 | 0.9976 | 0.9316 | 0.9875 | 0.9588 | 0.9984 |
| | 0.9326-0.9517 | 0.9679-0.9817 | 0.9518-0.9645 | 0.9971-0.9982 | 0.9212-0.9412 | 0.9824-0.9922 | 0.9526-0.9645 | 0.9979-0.9988 |
| LFC | 0.9424 | 0.9368 | 0.9396 | 0.9948 | 0.9589 | 0.9152 | 0.9366 | 0.9950 |
| | 0.9321-0.9523 | 0.9262-0.9470 | 0.9319-0.9471 | 0.9937-0.9959 | 0.9501-0.9680 | 0.9032-0.9265 | 0.9289-0.9440 | 0.9939-0.9960 |
| Micro avg | 0.9610 | 0.9610 | 0.9610 | 0.9977 | 0.9580 | 0.9580 | 0.9580 | 0.9973 |
| | 0.9568-0.9650 | 0.9568-0.9650 | 0.9568-0.9650 | 0.9972-0.9981 | 0.9538-0.9623 | 0.9538-0.9623 | 0.9538-0.9623 | 0.9969-0.9978 |
| Macro avg | 0.9613 | 0.9609 | 0.9610 | 0.9974 | 0.9589 | 0.9579 | 0.9580 | 0.9975 |
| | 0.9571-0.9653 | 0.9568-0.9649 | 0.9569-0.9650 | 0.9969-0.9979 | 0.9549-0.9630 | 0.9536-0.9622 | 0.9537-0.9622 | 0.9971-0.9980 |
| Weighted avg | 0.9612 | 0.9610 | 0.9610 | 0.9974 | 0.9588 | 0.9580 | 0.9580 | 0.9975 |
| | 0.9572-0.9652 | 0.9568-0.9650 | 0.9569-0.9650 | 0.9969-0.9979 | 0.9548-0.9629 | 0.9538-0.9623 | 0.9538-0.9622 | 0.9971-0.9980 |

**Table S3. Performance metrics of unified fixed neuron models. Related to Figure 3.**

The one-vs-rest precision, recall, F1-score and AUC for each cell type and a group of neurons, and the overall metrics (micro, macro and weighted averages) are shown. The 95% CI is indicated below each value.
